# Resolving endogenous protein organization in cells with nanometer precision

**DOI:** 10.1101/2025.08.11.669713

**Authors:** Janna Eilts, Marvin Jungblut, Dominic A. Helmerich, Stefan Sachs, Christian Werner, Christian-Alexandru Bogaciu, Ali H. Shaib, Silvio O. Rizzoli, Philip Kollmannsberger, Sören Doose, Markus Sauer

## Abstract

Advances in super-resolution microscopy have outpaced molecular labeling methods, impeding nanometer-scale imaging of endogenous multiprotein complexes in cells. To overcome these limitations, we developed an expansion microscopy (ExM) approach based on double-homogenized hydrogels that enables *direct* stochastic optical reconstruction microscopy (*d*STORM) of 7-8-fold expanded immunolabeled samples. The resulting ∼4-fold increase in effective immunolabeling density resolved the 8 nm distance between neighboring α-tubulin molecules in microtubules and the polyhedral lattice in clathrin-coated pits with nanometer precision in cells. Two-color Ex-*d*STORM further revealed the molecular organization of RIM scaffolding protein and Munc13-1, an essential synaptic vesicle priming protein, in ring-like structures with diameters of 40-45 nm at the presynapse in hippocampal neurons. Our results demonstrate that Ex-*d*STORM resolves the molecular organization of endogenous multiprotein complexes with nanometer spatial resolution in genetically unmodified cells. Thus, it provides a versatile method for the investigation of molecular protein distributions in their physiologically relevant context.

**One-Sentence Summary:** *The optimized combination of dSTORM and double TREx ExM resolves previously unresolvable protein assemblies in cells*

## Main Text

Latest single-molecule localization microscopy methods demonstrated superior localization precisions and spatial resolutions in the nanometer range on isolated molecules (*1-4*). Similarly, expansion microscopy (ExM) combined with fluorescence intensity fluctuation analysis could resolve the shape and conformation of isolated proteins (*5,6*). However, these methods can currently not reliably image protein complexes and arrangements with molecular resolution in cells, respectively. This limitation is mainly caused by inefficient labeling with fluorescent probes such as IgG antibodies. But also, smaller probes such as fluorescent proteins and nanobodies as well as chemical tags impede stoichiometric labeling of multiprotein complexes and do not permit the labeling densities required to achieve molecular resolution imaging in cells.

A very promising method for high-density labeling of genetically unmodified proteins in cells with minimal linkage error represents post-expansion immunolabeling because dense protein structures are decrowded and epitopes better accessible by nanometer-sized probes after expansion (*7-9*). Particularly, in combination with single-molecule localization microscopy, expansion microscopy should thus enable fluorescence imaging of dense protein structures in cells with nanometer spatial resolution. Results reported about single-molecule localization microscopy of expanded samples were motivating but disclosed also problems that must be solved to unleash the full potential of the methodology (*10-12*). Among these problems are lower expansion factors and inhomogeneous expansion achieved when trying to preserve protein epitopes for post-expansion immunolabeling (*7,13*). Furthermore, addition of a photoswitching buffer as required for *direct* stochastic optical reconstruction microscopy (*d*STORM) to a swellable polyelectrolyte hydrogel with hydrophilic ionic side groups results in shrinking of the gel (*14*). Albeit shrinking can be reduced by re-embedding of charged expanded hydrogels in an uncharged polyacrylamide gel (*10,15,16*), the achieved expansion factors were only moderate (*10-12*), and carbocyanine dyes such as Alexa Fluor 647 (AF647), which are the most suitable dyes for *d*STORM, are efficiently destroyed during re-embedding (*13,17*). Therefore, sub-10 nm imaging by ExM in cells remains challenging.

In this study, we overcome these previous limitations by using a refined post-expansion immunolabeling and re-embedding protocol that uses two Ten-fold Robust Expansion Microscopy (TREx) steps (*18*) combined with re-embedding in a neutral hydrogel to achieve 7-8-fold expansion of dense cellular structures and efficient localization of carbocyanine dye labeled proteins. ExM combined with *d*STORM (Ex-*d*STORM) allowed us to resolve cellular multiprotein assemblies including the periodicity of microtubules (*19,20*), the polyhedral lattice in clathrin-coated pits (*21,22*), the molecular organization of the nuclear pore complex (NPC) (*23,24*), and presynaptic proteins of the vesicle docking machinery with unprecedented resolution in genetically unmodified cells (*25-27*).

### Ex-dSTORM in double-homogenized hydrogels

A broadly applicable nanoscopy method should allow us to resolve the molecular architecture of endogenous proteins and complexes in their physiologically relevant context, i.e. in genetically unperturbed cells, while relying on accessible and widely adopted labeling strategies. Immunolabeling with antibodies best fulfills these criteria, as antibodies are well established, sufficiently sensitive, and available for most target proteins. Post-expansion immunolabeling offers additional advantages: it enhances epitope accessibility and reduces the effective linkage error in proportion to the expansion factor (*10*). However, since milder homogenization is required to ensure epitope survival for post-expansion labeling with antibodies, the resulting expansion factors are small and often inhomogeneous (*7,13,18,28*). This becomes particularly problematic when strong fixation or anchoring methods are used in combination with higher expansion factor protocols. Alternatively, some protocols avoid crosslinking by aldehyde fixation to enable homogenization by denaturation (*7,28,29*).

To maximize protein retention, epitope survival, and efficient anchoring of proteins into hydrogels we developed a two-step post-expansion immunolabeling method that uses standard fixation with aldehydes and protein denaturation to preserve epitopes during the initial expansion step. After immunolabeling with primary antibodies, the sample is enzymatically digested by proteinase K treatment followed by immunolabeling with secondary antibodies (Fig. 1). In the first expansion round, cells are fixed with glutaraldehyde (GA) or formaldehyde (FA) and anchored into the hydrogel using GA or a combination of FA and acrylamide (FA+AA). After gelation with TREx monomer solution (18), proteins anchored in the hydrogel are denatured with sodium dodecyl sulfate (SDS) and dithiothreitol (DTT) at 98°C to ensure epitope preservation. During immunolabeling with primary antibodies in phosphate buffered saline (PBS), the hydrogel remains ∼3-fold expanded, which increases the distance between adjacent epitopes and promotes higher labeling densities with IgG antibodies. Next, the first hydrogel is crosslinked and re-embedded into a second TREx gel, causing the gel to slightly shrink to a ∼2.5-fold expanded state. Anchoring with glutaraldehyde ensures that the primary antibodies are efficiently linked into the second hydrogel matrix. After proteinase K digestion for 45-120 min at 37°C, the sample is immunostained with fluorophore-labeled secondary antibodies. Finally, the gel is fully expanded (8-9x) and sliced with a razorblade to facilitate diffusion of neutral (non-expanding) gel monomer solution required for stabilization of the expanded sample for *d*STORM imaging (Fig. 1).

**Fig. 1.**
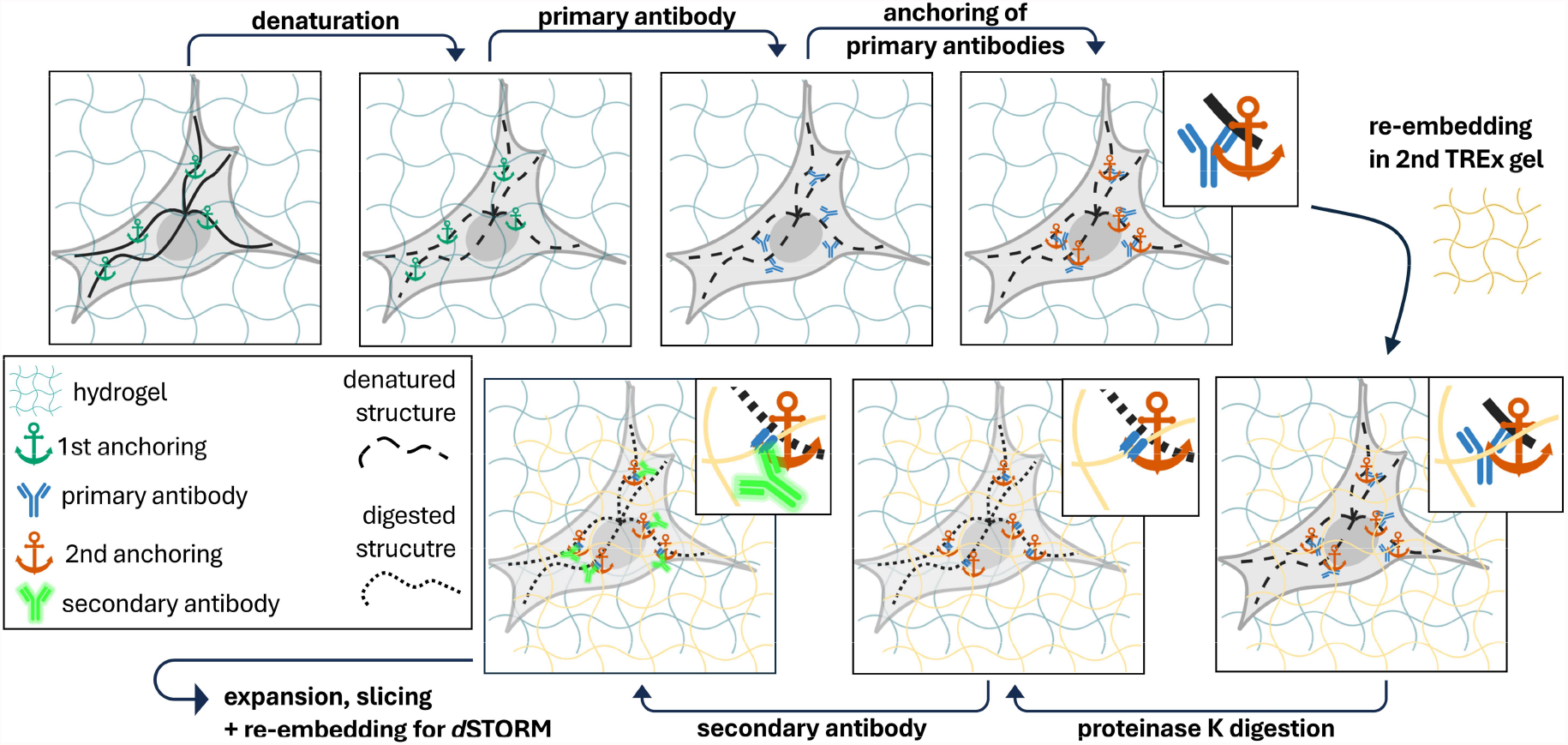
*Double* TREx (*d*TREx) expansion protocol with improved homogenization. Fixed cells are anchored into the hydrogel matrix, followed by a gentle homogenization step through denaturation. This initial, mild treatment allows for efficient labeling of proteins with primary antibodies, which are subsequently incorporated into a second TREx gel. Once the second gel has polymerized, a more rigorous homogenization is carried out using proteinase K digestion. Finally, secondary antibodies are applied to label primary antibody fragments that are anchored into the second gel and the sample is fully expanded. To prepare the gel for *d*STORM imaging, it is sectioned in thin slices and re-embedded into a neutral gel matrix. *Created with Biorender*.*com*.

We applied the *double* TREx protocol (*d*TREx) on microtubules and clathrin-coated pits in GA-fixed and anchored COS-7 cells and compared the performance with standard TREx post-expansion immunolabeling and *d*TREx omitting the proteinase K step, i.e. using only denaturation for homogenization. The resulting Airyscan microscopy images clearly showed that TREx as well as *d*TREx with denaturation alone result in ruptured expanded microtubules and poorly expanded clathrin-coated pits lacking detailed ultrastructural features about the clathrin coat. In contrast, microtubules expanded by *d*TREx using additional digestion with proteinase K were imaged as continuous filaments (fig. S1). Double homogenization with *d*TREx resulted in expansion factors of the hydrogel (the macroscopic expansion factor) of 7.6 ± 0.3 (s.d.) and 8.4 ± 0.6 (s.d.) using proteinase K digestion at 37°C for 45 min and 2 h, respectively. Interestingly, *d*TREx using denaturation only showed a similar macroscopic expansion factor of 7.6 ± 0.4 (s.d.) (fig. S1e and S2). On the other hand, the microscopic expansion factors determined from clathrin-coated pits and microtubule diameters showed that double homogenization using *d*TREx results in ∼2.5-fold higher expansion (fig. S1f,g), indicating that 8-9-fold expansion of multiprotein complexes requires an additional proteinase K digestion step. These findings underscore the importance of using a balanced protocol that ensures efficient and homogeneous expansion of multiprotein complexes in cells but concurrently preserves protein epitopes for post-expansion immunolabeling.

Having established *d*TREx for homogeneous 8-9-fold expansion of multiprotein complexes in cells, we next focused our efforts on optimizing re-embedding for *d*STORM imaging with carbocyanine dyes such as AF647. Unfortunately, carbocyanine dyes do not survive the gelation step, most probably due to a radical attack at one of the conjugated double-bonds during polymerization. Primary antibodies labeled with secondary AF647-antibodies only retain less than 10% of their initial brightness after gelation and digestion (*13*). Therefore, we tested the use of lower ammonium persulfate (APS) and tetramethylethylenediamine (TEMED) concentrations during re-embedding in the neutral gel to improve signal retention of AF647-labeled secondary antibodies. By titrating the radical starter and catalysator concentration we observed improved survival of AF647 in ensemble experiments with decreasing APS/TEMED concentration (fig. S3a). Re-embedding of *d*TREx gels with an APS/TEMED concentration of 0.025% resulted in only 10% hydrogel shrinking and ∼60% signal retention comparing the fluorescence intensity of identical microtubule filaments in the sample before and after re-embedding (fig. S3b-e). Accordingly, the final expansion factors are 10% smaller. Considering that secondary antibodies are labeled with an average degree of labeling (DOL) of ∼3, each antibody should be localized in *d*STORM experiments of expanded samples with high probability using the optimized re-embedding protocol. To avoid chemical destruction of carbocyanine dyes staining with the secondary antibody can be performed after re-embedding. However, while the measured fluorescence intensities of microtubules labeled either before or after re-embedding with secondary AF647-antibody showed similar fluorescence intensities, the background signal was slightly higher for samples labeled post-re-embedding in the neutral gel (fig. S4). Therefore, we labeled the samples before the re-embedding step in all following experiments.

### *d*TREx with post-expansion immunolabeling resolves the 8 nm distance between neighboring α-tubulin molecules

To test the performance of *d*TREx expansion and Ex-*d*STORM imaging we used microtubules as protein structure that is frequently used as cellular reference in super-resolution and expansion microscopy (*30,31*). Microtubules are polar cytoskeletal polymers that play essential roles in intracellular transport, cell division, and the spatial organization of the cytoplasm. Microtubules are assembled from α,ß-tubulin heterodimers, which stack head-to-tail into polar protofilaments with a periodicity of 8 nm. In most eukaryotic cells, thirteen protofilaments associate laterally to generate a hollow, cylindrical tube with an outer diameter of ∼25 nm. This conserved 13-protofilament architecture underlies the mechanical rigidity and dynamic behavior that characterize microtubules (*19,32*). The lateral arrangement of protofilaments within the microtubule wall predominantly follows a B-lattice configuration (*33*). In this geometry, α-tubulin subunits laterally contact α-tubulin, and β-tubulin contacts β-tubulin in neighboring protofilaments, preserving homotypic lateral interactions across most of the cylinder. However, because the helical pitch of tubulin polymerization does not perfectly match the geometry of a closed 13-protofilament tube, this regular B-lattice arrangement cannot be maintained continuously. As a result, a single discontinuity, termed the seam, forms along the microtubule wall. At this seam, lateral contacts adopt an A-lattice configuration, in which α-tubulin laterally interacts with β-tubulin with an offset of 3 tubulin monomers due to the helical pitch. Thus, canonical 13-protofilament microtubules are characterized by a largely uniform B-lattice organization interrupted by a single A-lattice seam. This structural feature is thought to influence microtubule stability, lattice plasticity, and interactions with microtubule-associated proteins, and represents a defining element of microtubule molecular architecture (*19,32-34*).

After immunolabeling microtubules exhibit an outer diameter of 60 nm accounting for a linkage error of 17.5 nm defined by the primary and secondary antibody (*32*). Although the periodic structure of microtubules is particularly suited as a reference for advanced super-resolution microscopy technologies (*1-4*), the limited labeling density along microtubules achieved when using IgG antibodies with a size of 10-15 nm (*32*) did so far not allow resolving the 8 nm distance between neighboring tubulin molecules. According to information theory, the required density of fluorescent probes must be sufficiently high to satisfy the Nyquist–Shannon sampling theorem (*35*). At its most basic level, the theorem states that the mean distance between neighboring localized fluorophores (the sampling interval) must be at least twice as fine as the desired resolution. With 13 protofilaments per ring, 1 µm of microtubule comprises 1,625 α-tubulins, of which only a few percent are detected in *d*STORM experiments of unexpanded microtubules labeled for α-tubulin with primary and secondary antibody, which is insufficient to resolve molecular details (fig. S5a). On the other hand, if microtubules are labeled with primary anti-α-tubulin antibodies after the first TREx expansion step, we detected 3 to 5 times as many localizations in *d*STORM experiments of fully *d*TREx expanded and re-embedded microtubules demonstrating that post-expansion immunolabeling achieves substantially higher labeling densities (fig. S5b). For microtubule expansion we used 2 h proteinase K digestion resulting in a final expansion factor of ∼8.0 (fig. S2) taking 10% shrinking during re-embedding in the neutral hydrogel into account.

The resulting 3D Ex-*d*STORM images of post-expansion immunolabeled α-tubulin in GA fixed and anchored COS-7 cells clearly indicate the presence of a periodic structure (Fig. 2a-c and fig. S6). However, since microtubules in B-lattice arrangement consist of typically 13 protofilaments with an offset of 0.923 nm between neighboring protofilaments, the periodicity cannot be easily identified by looking at entire stochastically labeled microtubules, where all axial distances occur, not only the 8 nm spacing along a single protofilament (fig. S7) (*33*). Only when examining the x,y projection of localization clusters detected by 3D *d*STORM along selected short filament areas, 8 nm distances between neighboring clusters can be visualized (Fig. 2b-e and fig. S6).

**Figure 2.**
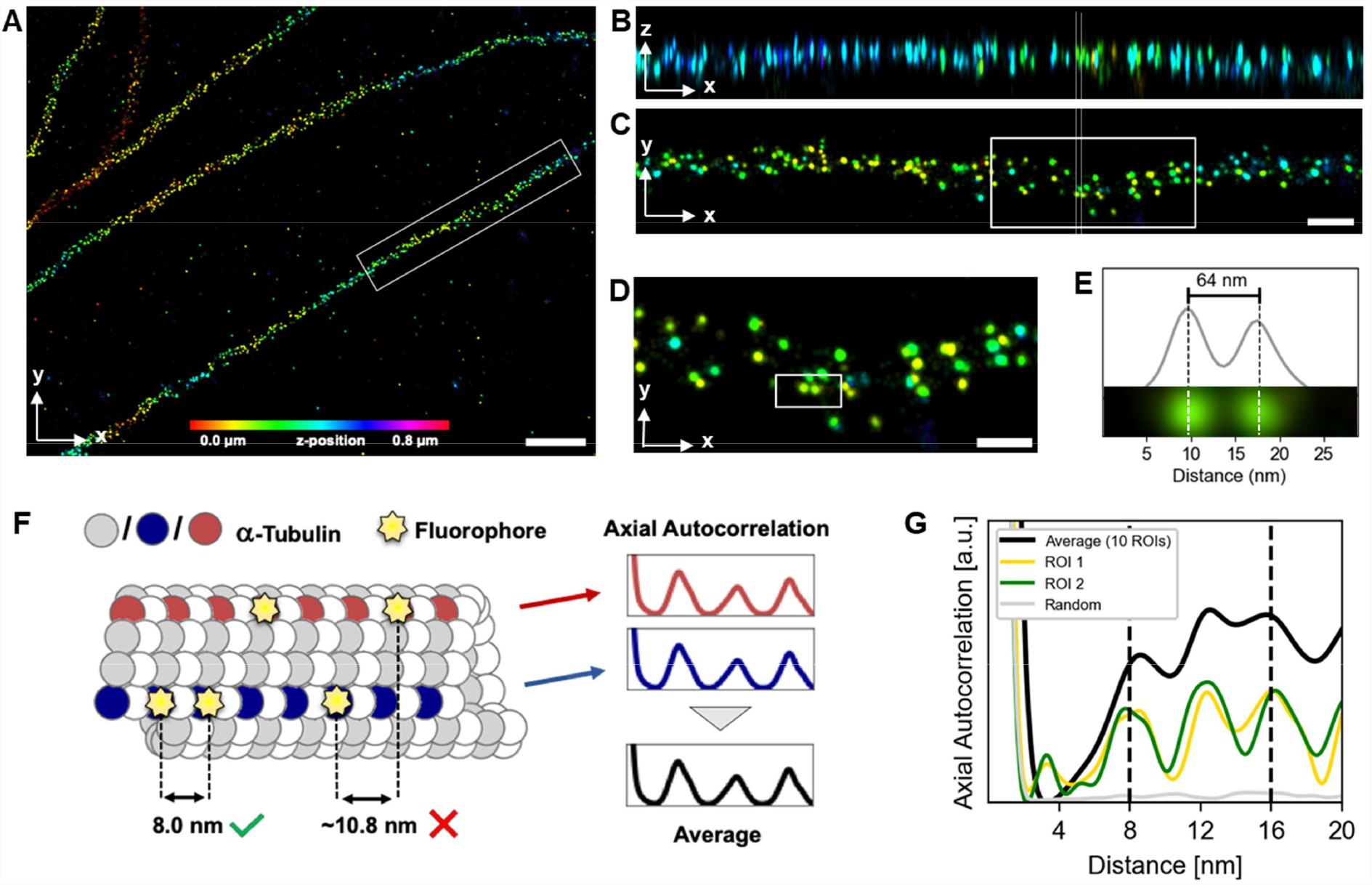
Ex-*d*STORM of microtubules reveals the 8 nm tubulin lattice. **(A)** Representative 3D Ex-*d*STORM image of a COS-7 cell using *d*TREx post-immunostaining for α-tubulin (AF647). **(B**,**C)** Corresponding xz- and xy-views of the region marked in (A). White dashed lines indicate a separation of 64 nm (8 nm unexpanded). **(D)** Zoomed-in view of the region marked in (C). **(E)** Zoomed-in view of region marked in (D), with intensity profile superimposed. The two signals are separated by 64 nm (8 nm unexpanded), corresponding to neighboring α-tubulin molecules on the same protofilament. **(F)** Schematic of a microtubule in B-lattice configuration where α-tubulin was stochastically labelled (yellow). Due to the lateral offset between protofilaments, the 8 nm spacing can only be directly observed along the same protofilament, while axial autocorrelation averaged over all protofilaments reveals the lattice spacing. **(G)** Autocorrelation function of two individual microtubules (green) and averaged over 10 microtubules (black). The same number of clusters randomly placed across the filament shows no detectable periodicity (gray) (fig. S8). The intensity profile in (E) and the autocorrelation analysis (G) were corrected for the final expansion factor of ∼8.0 after *d*TREx and re-embedding in a neutral hydrogel. Scale bars (expanded), A, 2 µm; B,C, 500 nm; D, 300 nm. Pixel size, 5 nm.

To unequivocally prove that Ex-*d*STORM after *d*TREx can resolve 8 nm separated immunolabeled proteins in cells, we identified the localization clusters of individual α-tubulins, assigned them to their respective protofilaments and analyzed the distances between them along individual filaments fig. S9). While the labeling density is not sufficient to see many neighboring α-tubulins, the tubulin lattice can be observed as peaks in the autocorrelation function of the signal along each protofilament at multiples of the lattice spacing (Fig. 2f,g and fig. S6). Here, the autocorrelation function clearly shows peaks at ∼4, 8, 12, and 16 nm as expected for microtubules in B-lattice configuration (Fig. 2f,g). The peaks at ∼4 nm and ∼12 nm can be explained by the α-tubulin signals along the seam of the B-lattice considering a single-seam of A-lattice contacts (*36*), which should give rise to the appearance of a 4 nm displaced, i.e. a 4 nm and 12 nm periodicity along the seam, respectively (fig. S7). Ex-*d*STORM experiments performed on ∼3.2-fold expanded cells cannot resolve the 8 nm tubulin lattice (fig. S10).

### Ex-dSTORM visualizes molecular details of clathrin-coated pits

Another popular reference structure for super-resolution microscopy is clathrin-coated pits (CCPs) (*11*). CCPs are curved to spherical, cage-like structures with a size of 50-200 nm located on the plasma membrane responsible for receptor-mediated endocytosis (*37,38*). Clathrin-mediated endocytosis involves more than 50 proteins and plays a key role in vesicular trafficking that transports a wide range of cargo molecules from the cell surface to the interior (*39*). The clathrin assembly unit is a trimer of three heavy chains and tightly associated clathrin light chains that form a “three-legged structure” termed triskelion with each leg measuring approximately 3 nm in thickness and 52 nm in length, ending in a globular “terminal domain” of 5 nm radius (Fig. 3a) (*40*). When assembled into a coat the legs of the triskelia interdigitate to form ordered lattices with variable ratios of predominantly pentagonal and hexagonal faces (Fig. 3b) (*21*).

**Figure 3.**
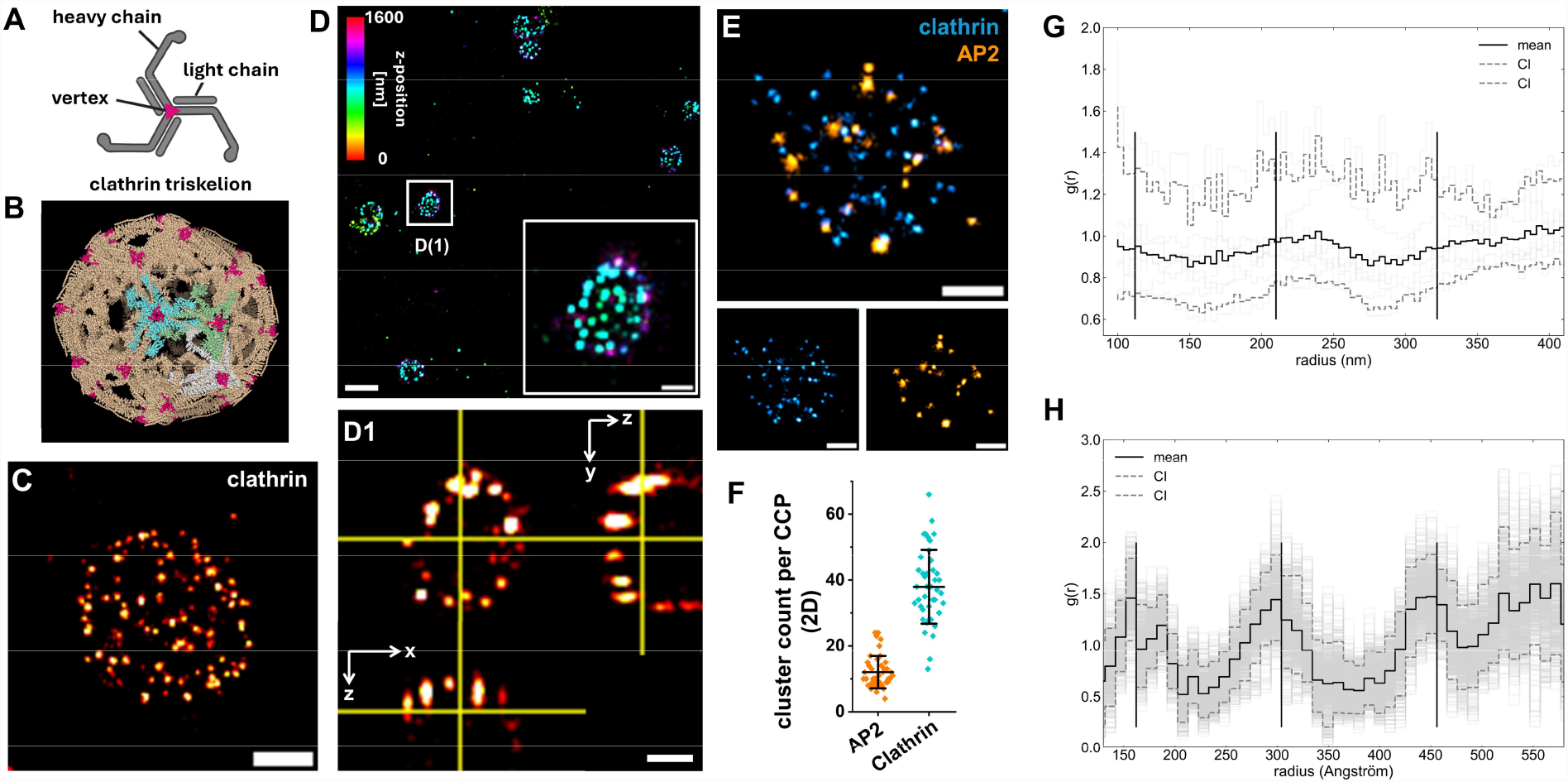
Ex-*d*STORM allows fluorescence imaging of CCPs with molecular resolution. **(A)** Schematic triskelion (created with Biorender.com). **(B)** Clathrin D6 coat reconstructed in Pymol from PDB structure 1XI4. Three exemplary triskelia are highlighted in cyan, green and white. Trimerization domains at the vertex of the triskelia are colored pink, representing binding sites for the used antibody. **(C)** Representative CCP visualized by Ex-*d*STORM using the *d*TREx procedure for enhanced homogenization and post-immunostaining for clathrin heavy chain (AF647). Cells were fixed and anchored with GA and proteinase K digestion was applied for 45 min at 37°C in the second TREx expansion step. **(D)** 3D Ex-*d*STORM image of CCPs. A magnified CCP is shown in the white box and as cross section (D1). **(E)** 2-color Ex-*d*STORM image of adaptor complex AP2 (AF647) and clathrin (CF568). **(F)** Localization clusters detected for AP2 and clathrin per CCP using DBSCAN. Mean values ± sd (AP2 = 12.0 ± 4.9, clathrin = 38.0 ± 11.2). Data from two independent experiments (n = 42). **(G)** Radial distribution function for all pairwise localization distances from the recorded *d*STORM localization data in selected regions of interest (n=8). Radial distribution functions are shown relative to those for localizations distributed under complete spatial randomness. Confidence intervals for 5% and 95% are shown (n=40). **(H)** Radial distribution functions for vertices of model structure shown in b (pdb: 1XI4). Radial distribution functions are generated and plotted with 5/95 % confidence intervals (n=100 random rotations). Scale bars, C,D1,E, 500 nm; D, 1 µm. Pixel size, c,e, 20 nm; D,D1, 35 nm. Scale bars are given for 7-fold expanded cells.

For visualization of CCPs by Ex-*d*STORM we immunolabeled the C-terminal region of clathrin heavy chain in GA fixed and anchored COS-7 cells. The C-terminal region of the clathrin heavy chain is located close to the vertex of the triskelion (*41*). After implementing the *d*TREx procedure with additional proteinase K digestion for 45 min at 37°C, gels were labeled with secondary antibodies and re-embedded in the neutral hydrogel. Ex-*d*STORM images clearly resolve the polyhedral lattice of expanded CCPs with an average size of 1 µm (Fig. 3c and fig. S11). The vertices of CCPs are visualized by clusters of localizations resulting from repetitive blinking of AF647-labeled secondary antibodies. 3D-Ex-*d*STORM images show the curvature of a just being formed CCP at the plasma membrane (Fig. 3d and fig. S12). Clathrin shells with average diameters of ∼80 nm contain about 35-40 triskelia and ∼20-25 heterotetrameric AP2 adaptors. AP2 adaptors link the clathrin coat and the membrane bilayer, and they are the principal cargo-recognition molecules (*41*). In our Ex-*d*STORM images we counted on average 12 AP2 and 38 triskelia assuming each localization cluster detected marks the vertex of a triskelion (Fig. 3e,f). This result confirms that our optimized *d*TREx and re-embedding protocol enables high density immunolabeling of endogenous proteins in cells and ensures efficient survival and localization of AF647-labeled antibodies.

Next, we determined the pairwise distances between all localizations from 2D-projected Ex-*d*STORM images of clathrin. The resulting radial distance distribution from all individual localizations of individual CCPs revealed peak positions around 100-150, 200-250, and 350-400 nm corresponding to distances of 14-21, 29-36, and 50-57 nm, respectively, assuming 7.0-fold expanded CCPs after re-embedding (Fig. 3g and fig. S13). The experimentally detected distribution is in good agreement with that for 2D-projected antibody epitope positions generated from the published D6 clathrin coat (pdb: 1XI4) which revealed peak distances of ∼16 nm, ∼30 nm and 46 nm (Fig. 3h and fig. S14). The peaks match the distance between adjacent vertices of the clathrin lattice, which have been measured by electron microscopy to 18-20 nm and 31-35 nm to the second-nearest vertex (*41,42*). The radial distance distribution of individual CCPs indicates some variation in peak positions and overall distribution possibly resulting from *d*STORM blinking effects and an estimated up to 10% expansion factor variation between different experiments. With an expansion factor of 7x, we determined an effective localization precision of ∼2.5 nm in lateral and 9 nm in axial direction in our CCP Ex-*d*STORM experiments even at an imaging depth of up to 100 µm. Here, the effective localization precision refers to the predicted localization precision from *d*STORM experiments divided by the expansion factor. The values demonstrate that *d*TREx with standard immunolabeling in combination with Ex-*d*STORM achieves a spatial resolution which is comparable to the size of a typical protein molecule in cells.

### Ex-dSTORM resolves endogenous NUP96 dimers in NPCs

To further demonstrate the performance of the method for resolving molecular details of cellular multiprotein complexes, we used the nuclear pore complex (NPC) as a well-characterized structural benchmark. NPCs rank among the largest macromolecular assemblies in cells and are composed of approximately 30 distinct proteins, termed nucleoporins (NUPs), which are present in multiple copies and arranged in an eightfold rotational symmetry (*23,24,43*). While super-resolution microscopy investigations of NPCs have been performed with genetically modified NUPs to improve the labeling efficiency as prerequisite for the resolution of nucleoporin dimers (*2,3,44,45*), we used COS7 cells and post-expansion immunolabeling to visualize endogenous NUP96 by Ex-*d*STORM. Cells were fixed with GA and FA and anchored with FA+AA. Processing according to the *d*TREx protocol included denaturation at 98°C and proteinase K digestion for 45 min at 37°C. After labeling with secondary antibodies gels were re-embedded in neutral hydrogel. NUP96 signals showed a mean peak-to-peak distance of 893 ± 48 nm (s.d.) translating into a microscopic expansion factor of 8.3x using the mean diameter of NPCs marked by NUP96 of 107 nm (Fig. 4a-c) (*45*). NUP96 is present in 32 copies per NPC and contributes to both the cytoplasmic and nucleoplasmic ring, with 16 copies in each ring. Within each ring, NUP96 is organized into eight dimers arranged at regular 12 nm intervals (*24,45*). Ex-*d*STORM allowed us to resolve the eightfold symmetry and endogenous NUP96 dimers separated by 11.9 ± 1.8 nm (mean ± s.d.) by standard immunolabeling (Fig. 4d-g).

**Figure 4.**
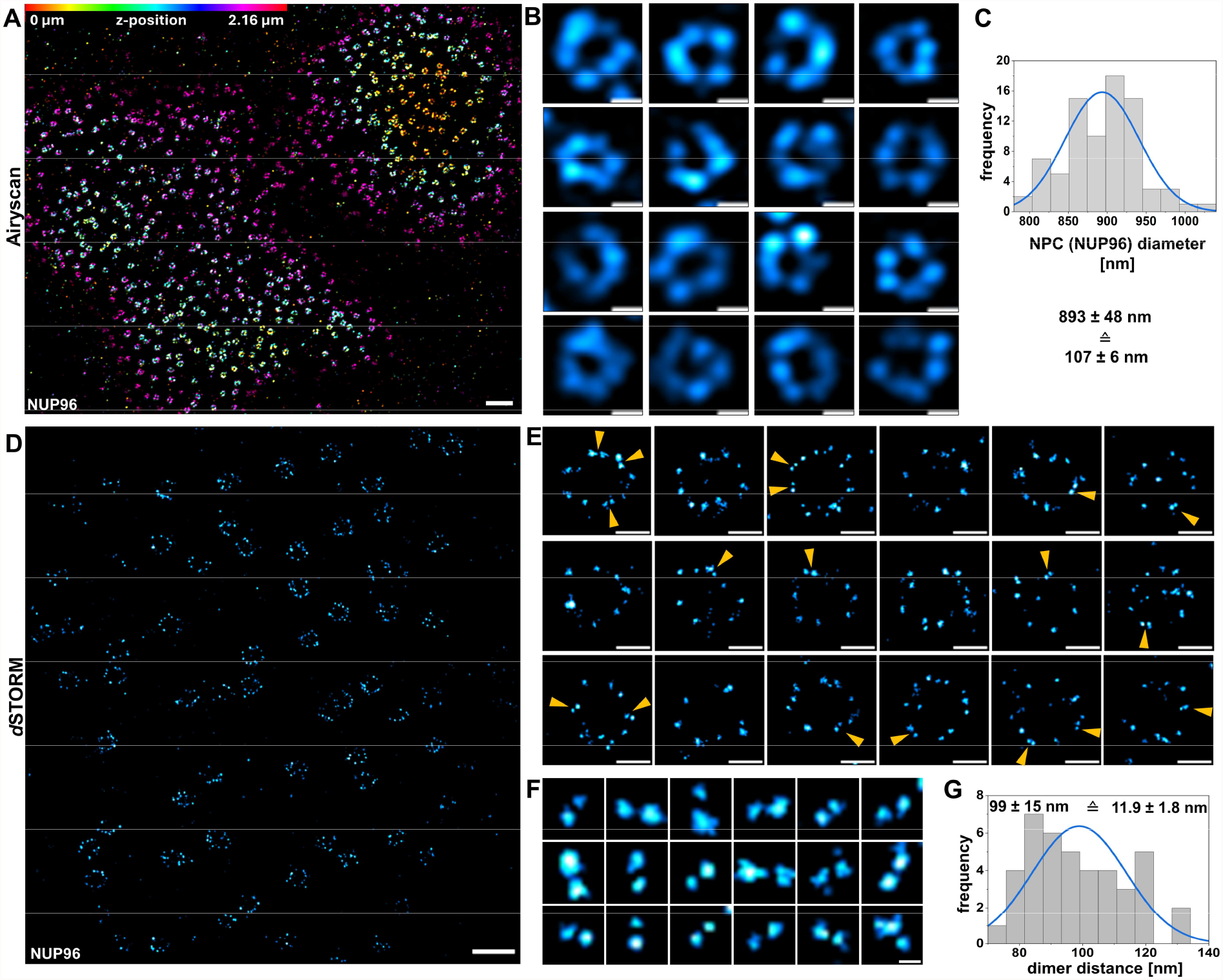
Ex-*d*STORM of endogenous NPCs in COS7 cells. **(A**,**B)** Airyscan images after re-embedding in the neutral hydrogel. Color-coded z-projection of nuclei immunostained for NUP96 (A). Magnified NPCs taken from single slices of Airyscan z-stacks (B). **(C)** Histogram of NPC diameters with normal distribution curve (blue) showing an average diameter of 893 ± 48 nm (mean ± s.d., n = 80 from 9 cells from one experiment) in Ex-*d*STORM images. This corresponds to ∼8.3-fold expansion after re-embedding considering the known mean diameter of 107 nm for NPCs marked by NUP96 (*45*). **(D-F)** *d*STORM images of re-embedded *d*TREx gels showing an overview (D) and magnified NPCs (E) immunostained for NUP96. Yellow arrows indicate NUP96 dimers. **(F)** Representative dimer signals used for distance measurements in (G). **(G)** Histogram of peak-to-peak distances of selected dimers with normal distribution curve (blue) showing an average distance of 99 ± 15 nm (mean ± s.d., n = 41) corresponding to 11.9 ± 1.8 nm considering the previously in (C) determined expansion factor of 8.3. Data from one experiment. Scale bars, A, 5 µm; B,E, 0.5 µm; D, 2 µm; F, 100 nm. Scale bars show 8.3x expanded dimensions after re-embedding in the neutral hydrogel.

### The presynaptic docking machinery is arranged in ring-like structures of different stoichiometry

Having established an efficient method for sub-10 nm fluorescence imaging in cells we reasoned that *d*TREx and Ex-*d*STORM can be used advantageously to resolve the molecular organization of Munc13-1 and Rab3-interacting molecule (RIM), two proteins that are important components of the vesicle fusion machinery in neurons at the presynaptic membrane (*46-48*). In central synapses the readily releasable pool typically consists of 5 to 10 synaptic vesicles (SVs) docked within 5 nm of the presynaptic plasma membrane (*25*). The release of neurotransmitters from SVs is remarkably fast requiring cooperation of multiple SNARE (soluble N-ehtylmaleimide-sensitive-factor attachment receptor) complexes – a key element of the process that mediates contact between SV and plasma membrane – to achieve this feat (*25,46,47*). Using cryo-electron tomography it has been shown that efficient binding of single SV requires a minimum of 6 copies of the chaperone Munc13-1 to initiate SV priming (*25,49*). Accordingly 6 copies of Munc13-1 can assemble into a closed hexagon with a diameter of ∼30 nm, which could, in cooperation with other presynaptic proteins of the docking machinery, form the template for SV capturing, priming, and release. Depending on the state of vesicle docking the orientation of Munc13-1 on the membrane changes and therefore the size of the hexagon might be variable (*25,50*).

The scaffolding protein RIM is closely interacting with Munc13-1 at presynaptic SV docking sites by binding via its Zinc-finger domain to the Munc13-1 C2A domain forming RIM-Munc13-1 heterodimers (*51*). Using immunolabeling and *d*STORM it was shown that both RIM and Munc13-1 form discrete nanoclusters at the presynapse that could represent the postulated template for SV docking (*52,53*). Achieving an improved immunolabeling efficiency and a 7-8-fold higher spatial resolution we hypothesized that our *d*TREx protocol in combination with Ex-*d*STORM can resolve these discrete nanoclusters and visualize the molecular organization of SV docking sites at the presynaptic membrane. Therefore, we immunolabeled Munc13-1 with AF647 and RIM1/2 (further referred to as RIM) with CF568 in hippocampal mouse neurons before re-embedding of *d*TREx-expanded hydrogels. The expansion factor was determined to ∼7.5 after re-embedding by comparing neurons immunolabeled for neurofilament-H before and after *d*TREx (fig. S15).

Since phorbol esters have been reported to increase neurotransmitter release by activating Munc13-1, hippocampal neurons were treated with 2 µM phorbol 12-myristate 13-acetate (PMA) for 30 min before fixation to increase the probability of detecting active vesicle fusion sites (*54*).

After fixation with FA and anchoring with FA+AA into the first TREx gel, samples were denatured at 98°C and labeled with primary antibodies. Further processing according to our *d*TREx approach (Fig. 1) included formation of the second TREx hydrogel and 45 min proteinase K digestion at 37°C before labeling with secondary antibodies and subsequent re-embedding into the neutral hydrogel. In the resulting 2-color Ex-*d*STORM images we focused on investigating frontal views of synapses to determine the molecular organization of Munc13-1 and RIM at docking sites at the presynaptic membrane (Fig. 5a and fig. S16). Indeed, using post-expansion immunolabeling in combination with two-color Ex-*d*STORM visualizes the organization of Munc13-1 and RIM in ring-like structures at active zones in hippocampal neurons under different conditions with an effective localization precision of ∼2.5 nm (Fig. 5a and fig. S17). To test spatial clustering, we computed Ripley’s h-function which confirmed the visual impression of clustering on various length scales. The h-function computed for simulated data with homogeneously distributed emitters which only exhibit *d*STORM-based repetitive blinking remains outside the 5-95% confidence interval of h-functions for all experimental datasets up to ∼100 nm (corresponding to ∼ 750 nm in the *d*TREx expanded hydrogel) (fig. S18).

**Figure 5.**
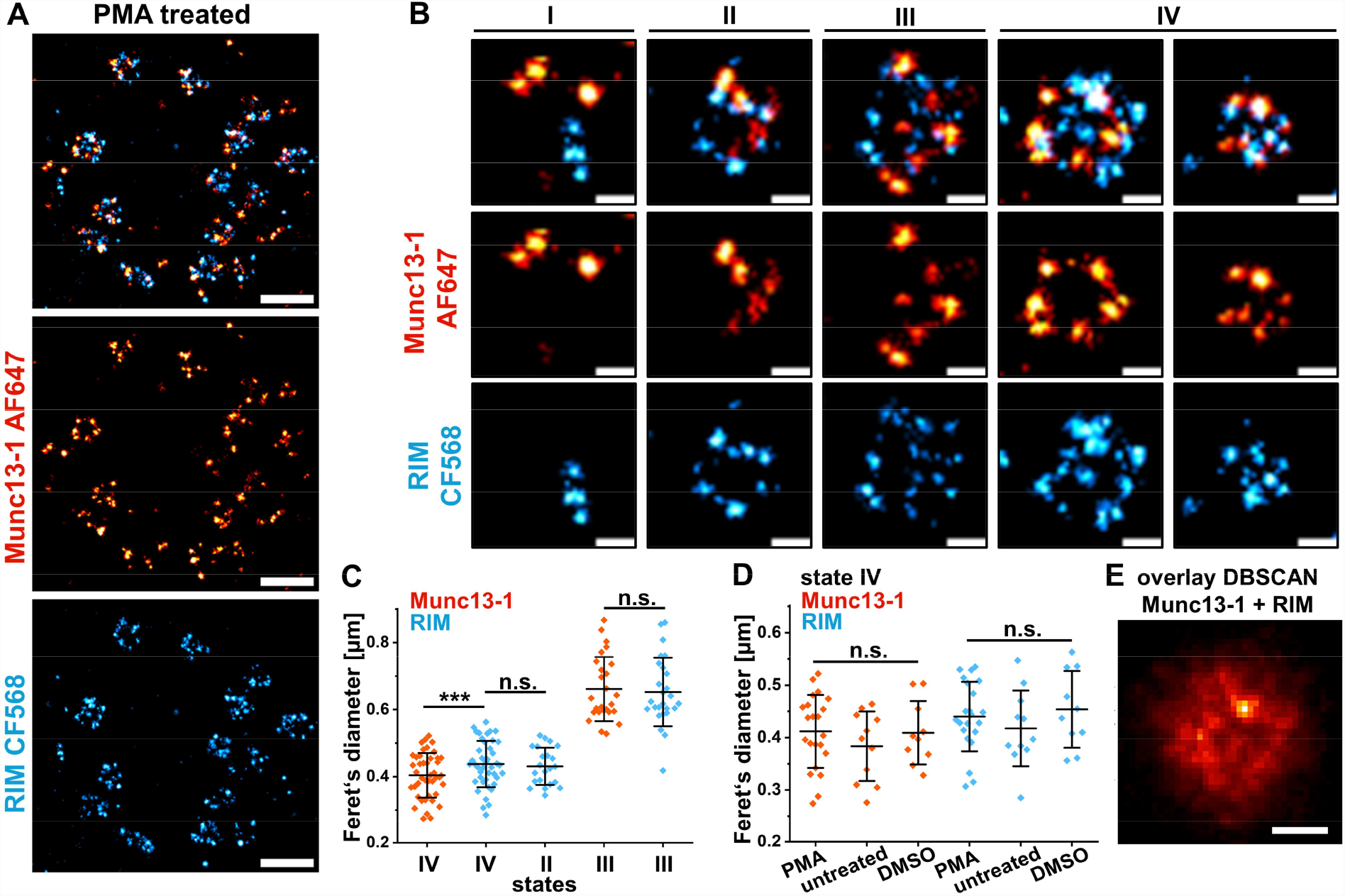
*d*TREx with two-color Ex-*d*STORM reveals the organization of Munc13-1 and RIM in hippocampal mouse neurons. **(A)** Two-color Ex-*d*STORM images showing a frontal view of a ∼7.5-fold expanded active zone in a hippocampal neuron. The sample was treated with 2µM phorbol 12-myristate 13-acetate (PMA) for 30 min before fixation to increase the probability of detecting active vesicle fusion sites (*54*). RIM (CF568, blue) and Munc13-1 (AF647, red) are organized in ring-like structures of different degrees that might represent SV docking sites. **(B)** Magnified Ex-*d*STORM images of individual Munc13-1 and RIM arrangements in active zones show regions of varying sub-structures from different synapses categorized in four different states. I: Munc13-1 and RIM unorganized. II: Only RIM shows ring-like arrangements. III: Munc13-1 and RIM are organized in substructures with a diameter > 500 nm. IV: Munc13-1 and RIM are both organized in ring-like structures with varying diameters < 500 nm. **(C)** Comparison of substructures exemplary shown in (B), fig S16 and S17. Feret’s diameter of structures determined using a polygon tool (fig. S19). In state IV ring-like structures of RIM exhibit a larger diameter (437 ± 69 nm) than Munc13-1 (404 ± 66 nm; n = 43) corresponding to ∼45 nm (RIM) and ∼40 nm (Munc13-1) diameters of unexpanded structures. The RIM diameter in state II (RIM = 431 ± 55 nm; n = 22) is identical to state IV. Munc13-1 and RIM diameters in state III are similar with 653 ± 102 nm (RIM) and 661 ± 96 nm (Munc13-1; n = 26) corresponding to diameters of ∼75 nm of unexpanded structures. Data was obtained from three independent experiments and neuronal cultures. Scatter dot graphs show mean ± s.d. P-values of paired sample t-test (for Munc13-1 versus RIM in state III/IV) and two sample t-test (for RIM state II versus state IV) shown as *** = p < 0.001 and n.s. (non-significant). **(D)** Munc13-1 and RIM diameters of ring-like structures determined in state IV under different experimental conditions (PMA treated, untreated and DMSO treated) show no significant (n.s.) differences (one-way ANOVA with post-hoc Tukey test). **e**, Overlay of localization clusters for Munc13-1 and RIM signals. Clusters were determined by DBSCAN and selected based on convex hull area and circularity, shifted by their centroid position and re-binned as overlay figure. The spatial distribution indicates the variety of substructures hiding the center hole that clearly appears in selected clusters. Scale bars, A, 1 µm; B,E, 200 nm. Scale bars show ∼ 7.5x expanded dimensions after re-embedding in the neutral hydrogel.

Visually two-color Ex-*d*STORM images indicate that Munc13-1 and RIM are organized in a variety of assemblies in active zones, most of them showing ring-like structures. To distinguish the different structures and classify them into four different states we used the following criteria (Fig. 5a,b, fig. S16 and S17): unorganized appearance of both signals (state I), only RIM exhibits ring-like arrangements (state II), Munc13-1 and RIM signals are organized in arrangements with diameters > 500 nm (state III), and both, Munc13-1 and RIM are organized in ring-like structures with varying diameter < 500 nm (state IV). While state I might represent Munc13-1 and RIM molecules that are in the process of forming SV docking sites, state II arrangements indicate that RIM forms ring-like structures in active zones more efficiently. Possibly preformed or faster assembled RIM rings recruit Munc13-1 to SV docking sites. State III might represent overlapping SV docking sites that cannot be clearly resolved under the applied experimental conditions. Finally, state IV shows smaller rings with diameters of ∼300-500 nm, probably representing established SV docking sites with varying stoichiometry of Munc13-1 and RIM (Fig. 5a-c, fig. S16 and S17). Here, the measured diameters of RIM structures were significantly larger than those of Munc13-1 structures (Fig. 5d).

Considering the expansion factor of 7.5x and the size of primary and secondary antibodies we determined average ring diameters for state IV structures in active zones of hippocampal neurons of ∼40 nm for Munc13-1 in accordance with previous reports (*25,55*) and ∼45 nm for RIM. Here it has to be considered that the measured ring diameters might be overestimated because we are labeling the N-terminus of Munc13-1 and RIM. Munc13-1 is a large protein with a size of ∼20 nm whereby the immunolabeled N-terminus directs away from the SV docking site, and the C-terminus is interacting with the synaptic vesicle (*56*). The sizes of the larger Munc13-1 and RIM clusters in state III did not show significant differences (Fig. 5c). In addition, we did not detect significant differences between the sizes of ring-like structures in state IV measured under different experimental conditions, i.e. PMA treated, untreated and DMSO (control) (Fig. 5d). Visual analysis of ring-like state IV structures under different experimental conditions indicates that we detect more ring-like assemblies after phorbol ester treatment which supports the idea that state IV represents functional SV docking sites (fig. S17).

Finally, we analyzed the number and size distribution of the ring-like structures by detecting localization clusters in the combined Munc13-1 and RIM signals with a DBSCAN algorithm and selecting cluster subsets based on convex hull area and circularity (figs. S19-S23) (*58,58*). It must be noted that this procedure is not capable of identifying all ring-like structures because there is too much overlap between individual localization clusters. However, the cluster selection procedure selects many well isolated clusters in an objective and reproducible way to compare cluster formation under different conditions. The overlay image of this cluster selection (Fig. 5e and fig. S21) and the calculated radial distribution function for signal clusters indicate identical length scales for Munc13-1 and RIM in and between all experimental conditions (fig. S22). Interestingly, applying only denaturation for homogenization (using a standard TREx gel) (18), does not lead to sufficient expansion of SV docking sites in active zones, highlighting the importance of the optimized *d*TREx protocol (fig. S24). Overall, our results demonstrate that active zones in hippocampal neurons exhibit strong differences in local protein density. In addition, SV docking sites show a large variation in composition and diameter, giving rise to the observation of different docking site arrangements.

## Discussion

Here, we introduce *d*TREx that enables a ∼4-fold higher labeling density by post-expansion immunolabeling with primary and secondary antibodies and thus allows in combination with Ex-*d*STORM to resolve the 8 nm distance of neighboring α-tubulin molecules in microtubules, fine details of the molecular architecture of clathrin-coated pits and nucleoporin dimers of nuclear pore complexes in genetically unmodified cells. Although conceptually not new, our refined *d*TREx protocol excels as it allows efficient expansion (8-9x before and 7-8x after re-embedding in the neutral hydrogel) combined with improved immunolabeling efficiency of endogenous proteins and reduced linkage errors. Optimization of the re-embedding conditions ensures that the majority of cyanine dyes (AF647) survive re-embedding and thus allow *d*STORM imaging in expanded gels. The combination of *d*TREx and Ex-*d*STORM can thus resolve the molecular organization of densely packed endogenous protein assemblies in cells.

Our results clearly show that the ability to resolve complex multiprotein structures in cells is not as much an issue of the localization precision provided by the super-resolution microscopy technique (*1-4*) as it is of the labeling density. The achieved effective localization precision of a few nanometers in combination with a ∼4-fold higher immunolabeling density enables researchers to uncover the nanoarchitecture of genetically unmodified cells using established immunolabeling and fluorescence imaging methods with so far unmatched structural resolution. The availability of imaging technologies that can visualize the nanoscopic organization of endogenous proteins in cells is important because transiently or constitutively overexpressed proteins can cause a multitude of problems especially in super-resolution microscopy experiments including protein aggregation and aberrant organelle morphology and generate misleading nanoscale structures and clustering artifacts (*59-61*). Our approach is broadly applicable since antibodies are available for efficient immunolabeling of most cellular proteins provided that the primary antibody binds well to epitopes of denatured proteins. For proteins that tend to aggregate during heat denaturation, this step may have to be adjusted using, e.g. a lower denaturation temperature.

Our results demonstrate that Ex-*d*STORM of *d*TREx expanded hippocampal neurons is ideally suited to unravel the molecular organization of synaptic proteins in crowded compartments. It allowed us to resolve the organization of Munc13-1 and RIM in active zones in ring-like assemblies of different diameters and geometries. The observation of a ring-like organization of RIM in active zones is supported by recent models that suggest that RIM is initially responsible for tethering of SVs to prepare them for membrane fusion (*62,62*). Different arrangements of the SV docking machinery in active zones might be well explained by the different states of vesicle docking, priming, and exocytosis (*25*). The diameters of the ring-like structures observed for Munc13-1 and RIM depend accordingly on the respective state of the individual docking site. In addition, exact diameters of these ring-like structures are difficult to determine given the uncertainties in linkage errors estimated for IgG antibodies after expansion (*9-13*). Nevertheless, the estimated ring diameters of ∼40 nm for Munc13-1 and ∼45 nm for RIM are comparable to protein density rings of ∼38 nm observed under primed vesicles by cryoelectron tomography (*55*). Altogether, our results demonstrate that the optimized *d*TREx protocol in combination with Ex-*d*STORM provides a versatile method to resolve previously unresolvable molecular details of the molecular organization of cells.

## ACKNOWLEDGEMENTS

The authors thank E. Maier, and I. Simeonov for cell culture and technical support, and S. Rizzoli and F. Opazo for cooperation and their constant support of this project.

## Author contributions

J.E. designed, performed and analyzed ExM experiments. M.J. performed and analyzed ensemble measurements. D.A.H. assisted in biplane imaging and performed the 3D analysis. S.S. assisted in LineProfiler analysis. C.W. prepared hippocampal mouse neurons. C.-A-B., A.H.S. and S.O.R. prepared neuronal samples and assisted data interpretation. S.D. and P.K. analyzed Ex-*d*STORM data. J.E, S.D., and M.S. interpreted the data. M. S. conceived the concept, designed experiments and supervised the study. All authors reviewed and approved the manuscript.

## Data and Materials availability

The data that support the findings of this study will be provided by the corresponding author upon reasonable request.

## SUPPLEMENTARY MATERIALS

Materials and Methods

References and Notes (64-71)

