## Supplementary material for "Resolving endogenous protein organization in cells with nanometer precision": Material and Methods and Supplementary Figures


**The PDF file includes:**

**Materials and Methods**

**Figs. S1 to S24**

**References and Notes**

### Materials and Methods

**Reagents.** B-27™ Plus Supplement (no. A3582801), formaldehyde methanol-free (no. 28906), GlutaMAX™ (no. 35050038), 10x HBSS (no. 14060040), Neurobasal™ Plus (no. A3582901), proteinase K (no. AM2548), sodium dodecyl sulfate (SDS, AM9820), Triton™ X-100 (Triton, no. 28314), trypsin-EDTA (no. 25300054), Tween 20 (no. 28320) were purchased from Thermo Fisher Scientific. Acetic acid (no. A6283), acrylamide (AA, no. A4058), ammonium persulfate (APS, no. A7460), (3-aminopropyl)triethoxysilane (APTES, no. 440140), bovine serum albumin (BSA, no. A7030), cysteamine hydrochloride (MEA, no. M6250), dithiothreitol (DTT, no. 646563), DMEM/F12 medium (no. D8062), ethanol (no. 32205), ethylenediaminetetraacetic acid disodium salt dihydrate (EDTA, no. E1644), ethylene glycol-bis(2-aminoethylether)-N,N,N′,N′-tetraacetic acid (EGTA, no. 03777), fetal bovine serum (FBS, no. F7524), formaldehyde (FA, no. F8775), gentamicin (no. G1272), glucose (no. G7528), glucose oxidase (no. G2133), glutaraldehyde (GA, no. G5882), guanidine hydrochloride (no. 50933), HEPES (no. H0887), KOH (no. P1767), 2-(N-morpholino)ethanesulfonic acid (MES, no. M3671), $MgCl_2$ (no. 442615), NaCl (no. S7653), N,N′-methylenebisacrylamide (Bis, M1533), PBS (no. D8537), penicillin-streptomycin (no. P4333), poly-D-lysine (PDL, no. P6407) and N,N,N′,N′-tetramethylethylenediamine (no. T7024) were purchased from Merck. Catalase (no. 6025.1) was purchased from Carl Roth GmbH. Phorbol 12-myristate 13-acetate (PMA, Cay10008014-1) was purchased from Biomol GmbH.

**Antibodies.** Rabbit anti α-tubulin (Abcam, no. ab18251), mouse anti α-tubulin (Merck, T6199), rabbit anti clathrin heavy chain (Abcam, no. ab21679) and mouse anti AP2 (Proteintech, no. 68349-1-Ig) were used at a concentration of 10 µg/ml. The working concentrations for neuron antibodies rabbit anti Munc13-1 (Synaptic Systems, no. 126103) and guinea pig anti RIM1/2 (Synaptic Systems, no. 140 205) were 15 µg/ml. Chicken anti neurofilament-H (Biolegend, no. 822601) was applied at 19 µg/ml for pre-expansion staining and at 38 µg/ml for post-expansion staining. Rabbit anti NUP98-96 (Proteintech, no. 12329-1-AP) was applied at a concentration of 15 µg/ml. Secondary antibodies donkey anti rabbit AF647 (ThermoFisher, no. A-31573), donkey anti mouse AF647 (ThermoFisher, no. A31571), donkey anti guinea pig CF568 (Merck, no. SAB4600469) and goat anti rabbit CF568 (Merck, no. SAB4600085) were used at a concentration of 20 µg/ml. Donkey anti chicken (Jackson Immunoresearch, no. 703-005-155) was conjugated with ~7-molar excess of N-hydroxysuccinimidyl-ester-CF568 (Merck, no. SCJ4600027) using Zeba™ Spin Desalting Columns (Fisher-Scientific, no. 87766) according to the instruction manual. The conjugated antibody was applied at a concentration of 10 µg/ml (pre-expansion) and 20 µg/ml (post-expansion).

**Cell culture.** COS-7 cells were cultured at 37°C with 5 % $CO_2$ in DMEM/F12 medium supplemented with 10 % FBS and 1% penicillin-streptomycin.

**Preparation of primary hippocampal mouse neurons.** Neurons were isolated from E18 C57BL/6 mice, in accordance with approval from the Bavarian state authorities. Hippocampal tissue was enzymatically digested using 0.05% trypsin-EDTA at 37°C for 15 minutes, followed by two rinses in HBSS. The HBSS solution consisted of 10x HBSS, 250 µl gentamicin, 3.5 ml of 1 M HEPES, and was diluted to a total of 500 ml using distilled water. After digestion, the tissue was gently dissociated using pipettes with different pore sizes in a neurobasal culture medium. This medium was composed of 100 ml Neurobasal™ Plus, 1 ml GlutaMAX™, 50 µl gentamicin, and 2% B-27™ Plus Supplement. The dissociated neurons were seeded onto 12 mm coverslips that had been coated with PDL (0.1 mg/ml, incubated at room temperature for 1 hour and rinsed twice with distilled water). Each coverslip received 40,000 cells, which were then cultured in neurobasal medium at 37°C in a 5% $CO_2$ incubator for 21 days. Half of the medium was replaced once per week. Optionally, neurons were treated with 2 µM PMA or 0.2 % DMSO (solvent for PMA) for 30 min at 37°C immediately before fixation. To fix the neurons, they were treated with 4% methanol-free formaldehyde for 15 minutes at room temperature and washed three times with PBS.

***d*TREx with double homogenization.** COS-7 cells were fixed in two steps with cytoskeleton buffer (CB-buffer, containing 10 mM MES, 150 mM NaCl, 5 mM EGTA, 5 mM glucose and 5 mM $MgCl_2$, pH 6.1). First, cells were incubated with 0.3 % GA + 0.25 % Triton in CB buffer for 60 seconds at 37°C and immediately afterwards fixed with 2 % GA in CB-buffer for 10 min. For NPC visualization COS-7 cells were fixed first with 0.3 % GA and 0.25 % Triton for 60 seconds at 37°C and then directly with 2.4 % FA for 20 min. Primary hippocampal neurons (DIV21) were fixed with 4 % methanol-free formaldehyde (FA) for 15 min. After three washing steps COS7-cells were crosslinked with 0.25 % GA for 15 min and neurons with a 4 % FA + 30 % AA overnight at 37°C (*64*). Samples were washed with PBS twice and dipped in TREx monomer solution once before gelation (*65*). TREx monomer solution contained 1.1 M sodium acrylate, 2 M AA, 0.009 % Bis, 1x PBS, 0.15 % TEMED and 0.15 % APS. Coverslips were then flipped on a 60 µl drop of TREx monomer solution with the cell side down. Polymerization was performed on ice for 15 min and then at RT for 1.5 h. Gels were then homogenized in pre-heated denaturation buffer consisting of 200 mM SDS, 200 mM NaCl and 50 mM Tris (pH 8) for 1 h at 98°C. Right before this step 65 mM DTT was added to the buffer. Denaturation buffer was then removed thoroughly by washing 5 x with ~5 ml prewarmed (37°C) PBS on a rotating wheel. Primary antibodies were incubated at a concentration of 10-15 µg/ml in 5 % BSA iteratively overnight at 4 °C and again with fresh antibody solution for 3 h at 37°C. Unbound antibodies were removed by washing 3x 15 min with 0.1 % Tween 20 in PBS (PBST) on a rotating wheel. To enable an additional digestion step, primary antibodies were anchored into a second TREx gel. Therefor the partly expanded samples (~3 fold) were crosslinked with 0.25 % GA for 20 min and washed three times for 15 min with PBS. For re-embedding into a second TREx gel samples were incubated twice for 30 min in 500 µl TREx monomer solution containing 0.03% TEMED/APS. This caused the gels to shrink to a ~2.5-fold expanded state. The monomer solution was removed and the gel was placed between two coverslips and incubated for 1.5 h at 37°C in a $N_2$-filled humidified chamber. For more efficient homogenization - additive to denaturation - a digestion step was included. Therefor gels were incubated with 8 U/ml proteinase K in digestion buffer (50 mM Tris + 1 mM EDTA + 0.8 M guanidine HCl + 0.5 % Triton) for 45 min (CCPs, neuronal proteins and NPCs) or 2 h (microtubules) at 37°C. Gels were then thoroughly washed 5x for 15 min with cold 0.1 % PBST. Secondary antibodies were incubated at a concentration of 20 µg/ml in 5 % BSA overnight and again with fresh antibody solution for 3 h at 37°C. After 3x 15 min washing steps with 0.1 % PBST gels were fully expanded in $ddH_2O$ preferably overnight before imaging or re-embedding.

**Post-labeling procedure with denaturation only**. Gels were processed in the same way as described in the previous section (*d*TREx with double homogenization) until after the primary antibody incubation step. After washing unbound primary antibodies secondary antibodies were incubated at a concentration of 20 µg/ml in 5% BSA overnight at RT or 3 h at 37°C. Following 3x

15 min washing steps with 0.1 % PBST gels were fully expanded in $ddH_2O$ preferably overnight before imaging or re-embedding.

**Silanization of coverslips.** Gels were placed on silanized 24-mm round coverslips (thickness 1.5H, Carl Roth, PK26.2) for imaging and re-embedding for *d*STORM. Coverslips were previously cleaned via three 15 min washing steps with first $ddH_2O$, then 1M KOH and in the end 99% ethanol in an ultrasound bath. After drying, coverslips were incubated with silane solution consisting of 0.001 % APTES, 80 % ethanol and 0.02 % acetic acid in $ddH_2O$. Each coverslip was covered with ~250 µl of silane solution and left to completely evaporate under a fume hood. Coverslips were washed twice with 99 % ethanol, left to dry and stored at -20°C.

**Re-embedding for *d*STORM.** To prevent shrinking of expanded gels in salt-containing imaging buffer, gels were re-embedded into a neutral gel. Therefor we followed the previously published re-embedding method with some modifications (*66*). All incubation steps were conducted in 2 ml tubes on a spinning wheel. First the expanded gels were cut with a razor blade into thin (~0.5-1 mm) slices to facilitate diffusion of buffer into the gels. A crosslinking step with 0.25 % GA in $ddH_2O$ for 20 min was implemented, followed by 3x 15 min washing steps with $ddH_2O$. Gels were incubated in re-embedding solution (10 % AA + 0.15 % Bis in $ddH_2O$) containing 0.025 % TEMED and 0.025 % APS twice for 30 min. After removing the solution gels were placed on silanized 24 mm coverslips with the cell side down. Another coverslip was placed on top. Polymerization occurred for 1.5 h in a $N_2$-filled humidified chamber at 37-40°C.

**Airyscan imaging.** Airyscan images were captured using an LSM 900 microscope equipped with Airyscan 2 (Zeiss) in super-resolution (SR) imaging mode, utilizing a C-Apochromat 40×/1.2 numerical aperture (NA) water-immersion objective (Zeiss) for expanded gels. For expansion factor determination a Plan-Apochromat 63x/1.4 NA oil objective (Zeiss) was used for pre-expansion recordings and a Plan-Apochromat 20x/0.8 NA or Plan-Apochromat 10x/0.45 NA air objective for post-expansion recordings. Appropriate excitation wavelengths and filter settings for the dyes were chosen through the dye presets in ZEN 2 blue software (Zeiss, version 3.5). To compare fluorescence intensity of AF647 before and after re-embedding all acquisition parameters were kept the same. All images were processed using the standard strength mode for 3D Airyscan processing.

***d*STORM image acquisition.** Re-embedded gels were incubated in switching buffer for 45 min and immersed in fresh switching buffer immediately before imaging. Switching buffer consisted of 100 mM cysteamine hydrochloride (MEA) and oxygen scavenger system (5 % glucose, 11 U/ml glucose oxidase and 220 U/ml catalase) in 1x PBS (pH adjusted to pH 7.6). Imaging chambers were purged with Argon gas and sealed with parafilm. *d*STORM measurements were performed on an inverted fluorescence widefield microscope (Olympus, IX-71) equipped with a nosepiece stage (Olympus, IX2-NPS) and two EMCCD cameras (Andor iXon Ultra DU-897). Biplane 3D images were acquired with an 60x oil objective (Olympus, NA 1.45 PlanApo) and HILO (highly inclined and laminated optical sheet) illumination using 4-7 $kW/cm^2$ of an appropriate laser (Toptica, iBeamSmart 640-S_11598). For recording on two EMCCD cameras simultaneously, a two-channel image splitter (TwinCam, Cairn Research) equipped with a 50/50 beamsplitter (Cairn Research) was used and the cameras were synchronized by a pulse generator (DG535, Stanford Research Systems). Two identical bandpass filters (RazorEdge LP Edge Filter 647 RU, Semrock) were positioned in front of the two cameras. A dirchroic mirror (FF410/504/582/669, Chroma) was placed in front of the objective to separate excitation and emission light. In addition, a quarterwave plate (Thorlabs, SAQWP05M) was mounted for excitation with circular polarized light. Subsequently, 45000-60000 frames were acquired on both cameras with a frame rate of 50 Hz. A UV-laser pulse was added if required to return more fluorophores to the on-state. For calibration, 100 nm microspheres (Invitrogen, T7279) were prepared by incubating ~$10^5$ beads/ml in 1xPBS (Sigma-Aldrich, D8537-500ML) with 50 mM $MgCl_2$ (Sigma-Aldrich, M9272-500G) adjusted to pH 7.4, for 15 min using eight chambered cover glass systems with high performance cover glass (Cellvis, C8-1.5H-N), followed by three washing steps prior calibration experiments. The calibration measurements were performed by using a piezo scanner (Pifoc, Physik Instrumente) driven with a LVPZT servo controller (E-662, Physik Instrumente) to move the objective. For two color *d*STORM imaging an oil-immersion objective (APON 60×, numerical aperture 1.49; Olympus) was employed. Alexa Fluor 647 and and CF568 were excited by 5-6 kW/cm² of a 639 nm laser and 4-7 kW/cm² of a 561 nm laser (Genesis MX 639-1000 STM and Genesis MX 561-500 STM, Coherent). A dichroic mirror (FF410/504/582/669, Chroma) was used to separate excitation from emission light. A beamsplitter (630 DCXR customized, Chroma) projected emission light through two different bandpass filters (607/70 and 679/41 BrightLine series, Semrock) on two separate EMCCD cameras. 20000-30000 frames of the two channels were acquired sequentially in Hilo illumination with a frame rate of 50 Hz. Images were reconstructed in the open-source software rapidSTORM 3.3 (*67*) usually with a pixel size of 20 nm. For two-color alignment 0.2 µm fluorescent microspheres (Invitrogen™ , T14792) were imaged in each channel and an alignment matrix was created with the ImageJ plugin bUnwarpJ. The matrix was then applied to the reconstructed images.

**3D biplane analysis.** For extracting 3D information of biplane *d*STORM recordings, an intensity-based analysis routine was used. First, the measured image stacks were analyzed with rapidSTORM 3.3 (*67*). Therefore, the FWHM was set to 360 nm, the intensity threshold was set to 250 ADC and the fit window radius to 1100 nm. The corresponding localization files from both cameras were then analyzed further using a custom written python script to calculate the 3D intensity ratios. The calibration file as well as the sample files measured, were analyzed in the same way as described elsewhere in detail but focused on the rapidSTORM analyzed intensity values for calculating the intensity ratios (*68,69*). For visualizing 3D images, ImageJ (version 2.16.0/1.54g) and ThunderSTORM (version 1.3) were used.

**Ensemble measurement of AF647 survival under different gelation conditions.** Fluorophore stability under gelation-like conditions was assessed by monitoring time-dependent fluorescence intensity of Alexa Fluor 647 conjugated antibodies in solutions containing varying concentrations of APS and TEMED. Measurements were performed in quartz glass cuvettes using a Jasco FP-8350 spectrofluorometer. Excitation was set to the absorption maximum of Alexa Fluor 647, and emission intensity was recorded every 30 seconds. The temperature of the measurement chamber was adjusted to 37°C. Photobleaching was minimized by opening the excitation shutter only during acquisition intervals. For measurements a donkey anti-rabbit IgG conjugated with AF647 (ThermoFisher, A31573; degree of labeling ~5) was diluted to 1 µM in 1x PBS. APS and TEMED were then added to the antibody solution at final concentrations corresponding to three gelation protocols: TREx (0.15 %), re-embedding (0.05 %), and modified re-embedding (0.025 %). Measurements were started immediately after APS addition. Emission intensities were normalized to the first time point.

**Fluorescence intensity pre vs. post re-embedding.** In addition to ensemble measurements, Airyscan images of alpha-tubulin in COS-7 cells were used to analyze the fluorescence intensity of the fluorophore AF647. Using the *d*TREx protocol images of the same cell before and after re-embedding in the neutral gel with 0.025 % APS and 0.025 % TEMED were taken. ROIs of identical tubulin strands were selected, and an auto threshold ("Otsu") was applied. After converting the image to a mask, the outline of the mask ("Create Selection") was saved in the ROI manager. This ROI was then applied to the original image and the mean fluorescence intensity in this region was measured.

**Evaluation of timepoint for adding secondary antibodies.** The potential different effect of adding secondary antibodies (sAbs) before (pre) or after (post) re-embedding in the neutral gel was analyzed. Therefore, *d*TREx gels labeled for α-tubulin were labeled with identical secondary antibody concentrations and incubation times either pre or post re-embedding into the neutral gel. In gels with sAbs added post re-embedding washing with PBST was conducted more thoroughly (5x 1 h) with an additional overnight washing step with gentle agitation. In contrast. in gels with sAbs added before re-embedding washing with PBST was done as in the standard protocol (3x 15 min). Airyscan images were recorded with the same settings for both conditions and the fluorescence intensity of microtubule strands was evaluated. The microtubule structure in overview images was defined in the same way as described in the previous section ("Fluorescence intensity pre vs. post re-embedding"). Then the fluorescence intensity in the applied ROIs to the original images was measured. Further, three ROIs in each overview image were drawn to measure the background fluorescence intensity. The intensity was then normalized to the ROI area for comparison of the two different conditions.

**Expansion factor determination.** The expansion factor was calculated by comparing Airyscan images of identical areas in the samples before and after expansion. On the one hand landmarks (e.g. the distance between the same two clathrin particles) were measured (as shown in Ext. Data Fig 1e). In addition, the expansion factor for COS-7 cells (microtubules and CCPs) and neurons was determined using a custom written python script as described before (*70*). Initially, images of the identical structure in the unexpanded (imaged with 63x oil objective) and expanded state (imaged with 10x or 20x objective) were registered via a rigid similarity transformation, providing the structural expansion factor. Optionally, a gaussian blur was applied to expanded images to better match the resolution of the unexpanded image and facilitate the registration. Further, a non-rigid affine transformation was implemented which is used to calculate the difference to the similarity transformation, representing nonlinear distortions depicted in a distortion vector map. The same script was used to determine the loss of expansion during the re-embedding process for *d*STORM. For the NPC experiment in COS-7 cells the expansion factor was estimated by measuring the diameter of NPCs as the fixation was different from the fixation for microtubules and CCPs in COS-7 cells.

**CCP quantification.** To evaluate the expansion efficiency of different protocols the diameter of CCPs was measured using 3D Airyscan images and Fiji. The center of each individual CCP was selected by going through the z-stack. A line profile was drawn over the CCP and a k-plot was generated. The diameter was determined by measuring the peak-to-peak distance in the resulting plot. If only one peak was observable, the FWHM was measured.

**Cluster quantification of AP2 and Clathrin.** In 2-color Ex-*d*STORM images of AP2 and Clathrin, clusters were classified and counted using DBSCAN and a custom written python script using LOCAN. DBSCAN parameters were epsilon = 20, meaning all points that lie in maximum distance of 20 nm to the next point are grouped into a cluster. All points that are reachable from at least 3 points (minPts = 3) belong to the cluster core. If a point is only reachable by one other point it is still included in the cluster as a border point. Thresholds of 80 photons for the AF647 channel and 400 photons for CF568 channel were set. Radial distribution functions for Ex-*d*STORM localizations were carried out with python-scripts based on the LOCAN package.

**LineProfiler measurements.** For comparing the expansion efficiency of filamentous microtubule structures after undergoing four different expansion protocols, LineProfiler was used to determine the average diameter of microtubules (*71*). Airyscan images of microtubules were recorded using the same settings for all conditions. During data processing, the pixel size was set to 65 nm. To compensate for noise or incomplete data, the following parameters were applied: intensity threshold of 2, spline interpolation parameter of 3, and a blur radius of 20. These configurations affect only the positioning and orientation of the line profiles and do not alter the underlying image data. For each condition a total of 270-550 line profiles were fitted with a Gaussian function, and the full width at half maximum (FWHM) of the fits was used to determine the overall diameter of the microtubules. For graphical illustration the line profiles were normalized to [0,1], centered and overlayed.

**NPC quantification.** Sizes of NPCs immunostained for NUP96 in *d*TREx gels were evaluated by measuring their maximum diameter using a line profile and its peak-to-peak distance in Ex-*d*STORM images of 9 different nuclei. Furthermore, potential NUP96-dimers were selected in these images and their distance was measured with a line profile and its peak-to-peak distance.

**Statistical analysis.** All ExM-*d*STORM experiments were repeated at least three times (3 independent experiments), except for AP2/Clathrin two color measurements (2 independent experiments) and NPC measurements (one experiment) with comparable results. Data was analyzed using the software OriginPro (Version 2021b, OriginLab Corporation). Scatter dot plots as well as their descriptions in the text show the mean value ± standard deviation (s.d.). The number of objects analyzed are represented as single data points in the graphs. For testing normality the Kolmogorov-Smirnov test was conducted. Significant differences between the mean values of two groups were determined with a two-sample t-test. For two connected samples a paired sample t-test was conducted. Significant differences between the mean values of more than two groups were tested using a one-way ANOVA with subsequent Tukey-test. P-values illustrated as * $\triangleq p < 0.05$, ** $\triangleq p < 0.01$, *** $\triangleq p < 0.001$, **** $\triangleq p < 0.0001$ and ns ≙ non-significant.

**Analysis of Microtubule distances.** We manually selected straight microtubule segments between microtubule intersections, automatically determined their 3D centerline based on cluster coordinates, and analyzed the axial distances of localizations in angular segments. To determine the central axis of MTs, we first detected clusters of localizations using *hdbscan* and then calculated the average position of the 10 nearest neighbors of each cluster. These average positions lie near the central axis of the MT and were thus used as initial center points. For each cluster, we calculated the distance to the nearest center point, and all clusters further away than a distance of 250 nm from the nearest center point were discarded. The procedure was then repeated with the remaining clusters. The final axial centerline was determined by fitting a bicubic spline (k=3) through the detected centerline points.

**Axial coordinate system.** The centerline was used to define a microtubule-local coordinate system for all localizations and cluster centers. Each localization/cluster is defined by its axial position along the MT, its angle measured counter-clockwise relative to 12:00, and its radial distance from the MT axis.

**Distance analysis.** Due to the angular offset between the protofilaments constituting the microtubule, no periodicity along the axis can be detected when averaging over all angular positions, since all axial distances would be equally probable. However, when moving along the MT axis at a given angular position, tubulin dimers are spaced 8 nm apart, and thus a periodicity of multiples of 8 nm (and multiples of 4 nm around the seam) should be detectable. We therefore calculated the axial autocorrelation function of a Gaussian-smoothed histogram ($\sigma$=2.5 nm) of all cluster positions along the MT axis separately for 16 angle bins to detect the periodicity of the microtubule lattice, followed by averaging over all angle bins.

**Simulation.** To calibrate the analysis and to compare the results to theoretical expectations, we simulated MTs with $\alpha$-$\beta$ tubulin heterodimers stacked axially in 13 parallel protofilaments with a helical offset of 12 nm. A given fraction of $\alpha$-tubulin monomers was randomly selected as labelled with primary antibody. An additional uncertainty of 2 nm (xy) and 5 nm (z) was used to define the final positions of the detected cluster centers. The labeling efficiency was estimated by comparing the number of detected cluster centers to the number of $\alpha$-tubulin monomers per MT length. We then performed the same distance analysis as described above using the simulated emitter positions as cluster centers.

**Ex-*d*STORM data analysis for two-color experiments in neurons.** Ex-*d*STORM data from Munc13-1/RIM two color measurements resulted from three independent experiments for each condition respectively. Using the localizations determined with rapidSTORM 3.3 (as described above), we calculated and displayed Ripley's H-function, a normalized Ripley's K-function. Computation was carried out for each ROI without edge correction. The averaged H-function and a 5-95% confidence interval was computed from 100 simulated data sets with localizations distributed on identical ROIs with identical number of localizations in each ROI according to complete spatial randomness or a Neyman-Scott process. The Neyman-Scott clustering process resembles typical *d*STORM data. It has homogeneously distributed parent events with each parent having n offspring events, where n is Poisson distributed with mean 10, and with the offspring positions having a Gaussian offset with a standard deviation of 22 nm. The maximum of the H-function indicates a distance that is between cluster radius and diameter and thus provides an estimate for the average cluster size. Localization clusters that contain larger ring-like localization clusters were determined by an arbitrary but reproducible algorithm to compare cluster features between the various treatment conditions. Clusters were determined by combining all Munc13-1 and RIM localizations, running a DBSCAN algorithm with epsilon = 100 nm and minPoints = 20, and selecting those clusters with a convex hull area between 100,000 and 500,000 nm$^2$, a isoperimetric quotient larger than 0.85, and a radial distance between 150 and 200 nm. All analysis procedures were carried out with python-scripts based on the LOCAN package.

**Estimation of Munc13-1 and RIM diameters in active zones.** The sizes of RIM and Munc13-1 substructures in reconstructed Ex-*d*STORM images were evaluated in Fiji. The polygon tool was used to outline the outer border of the assembly of ring-like structures (fig. S19). Subsequently Feret's diameter was determined to obtain the maximum diameter of the structure. The average diameter of ring-like structures in expanded active zones of hippocampal neurons was determined to a mean of 403 nm (Munc13-1) and 437 nm (RIM) independent of the experimental conditions (Fig. 5e-d). Since proteins are immunolabeled with a primary antibody after the first expansion step of the *d*TREx protocol we added 12.5 nm as an average size value for a primary IgG antibody. The primary antibody is digested and ~3-fold expanded in the second expansion step resulting in a displacement of ~37.5 nm. In addition, we used the secondary IgG antibody (+12.5 nm), which is added to the sample during the second expansion step. Hence, a value of ~100 nm (50 nm × 2 for both sides of the ring-like arrangement) must be subtracted from the measured diameters. This results in corrected average diameters of state IV of 303 nm (Munc13-1) and 337 nm (RIM). Divided by the overall expansion factor of 7.5x we obtain average ring diameters of ~40 nm for Munc13-1 and ~45 nm for RIM rings in state IV independent of the experimental conditions.

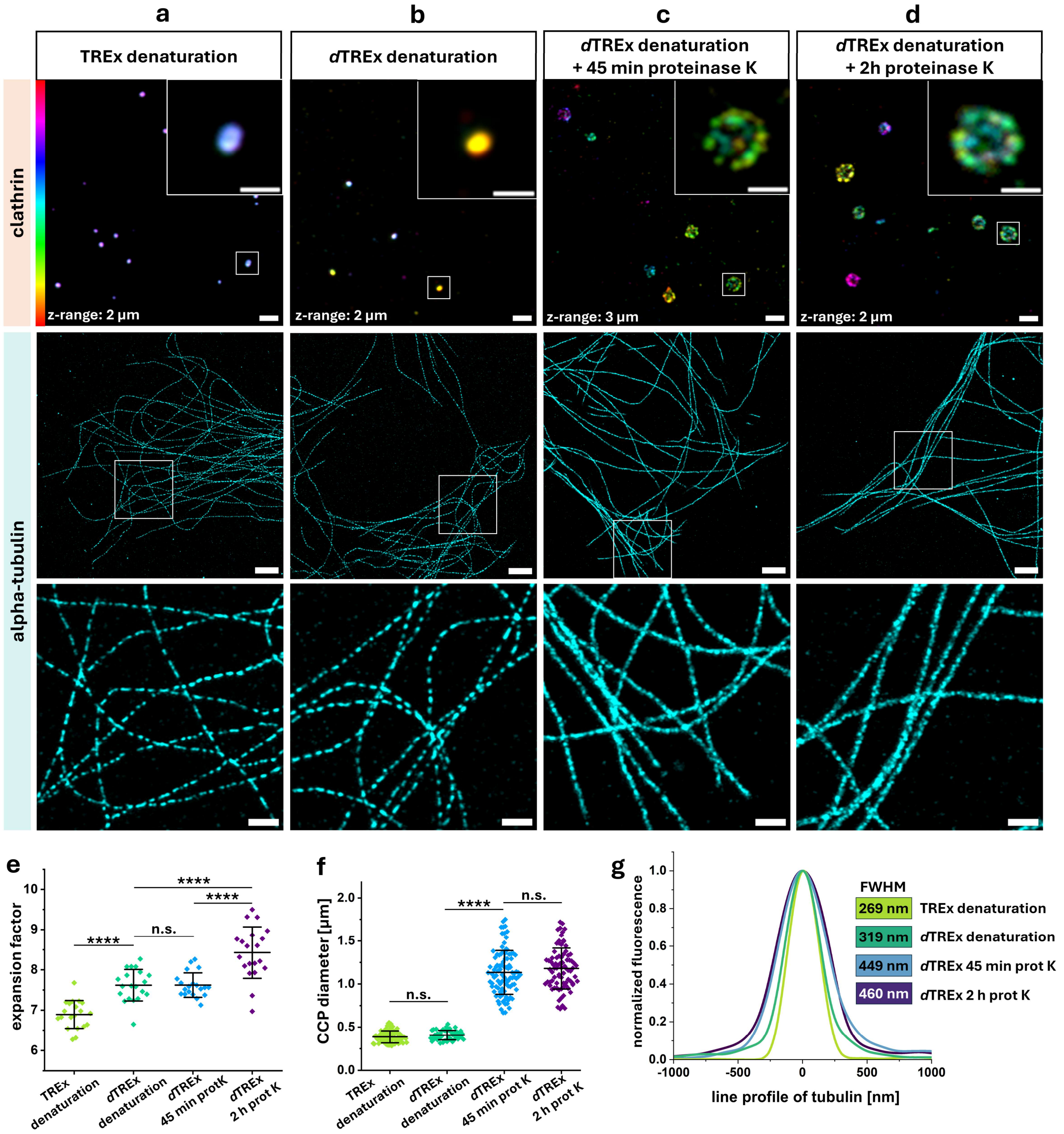


**Supplementary Fig. S1. Comparison of different post-immunolabeling approaches**. **(A-D)**, Airyscan fluorescence images of GA fixed and anchored COS-7 cells post-labeled after denaturation for clathrin heavy chain (upper row, scale bars 2 µm; magnified regions 1 µm) or alpha-tubulin (middle row, scale bars 10 µm and lower row for magnified regions, scale bars 3 µm) and processed according to the different protocols. Color code for clathrin images shows the z-range. **(A)** TREx with denaturation; **(B)** *d*TREx with denaturation; **(C)** *d*TREx with denaturation plus 45 min proteinase K digestion at 37°C and **(D)** *d*TREx with denaturation plus 2 h proteinase K digestion at 37°C. **(E)** Expansion factors of different protocols determined by measuring the distances between landmarks, i.e., the same two CCPs before and after expansion. Data from one experiment using the same monomer solution for all conditions. **(F)** CCP diameters determined using the different protocols. Data from three (TREx denaturation, n = 75; *d*TREx 45 min prot K, n = 94), two (*d*TREx 2 h proteinase K, n = 81) independent experiments and one experiment (*d*TREx denaturation, n = 50). **g**, Line profiles of single microtubule strands using different protocols measured by LineProfiler (*5*). The full width at half maximum (FWHM) of Gaussian fits was determined for each curve. Scatter dot plots show mean (line) ± s.d. (whiskers) and single data points (dots). P-values of one-way ANOVA with post-hoc Tukey-test are illustrated as **** ≙ $p < 0.0001$ and ns ≙ $p > 0.05$ (non-significant).

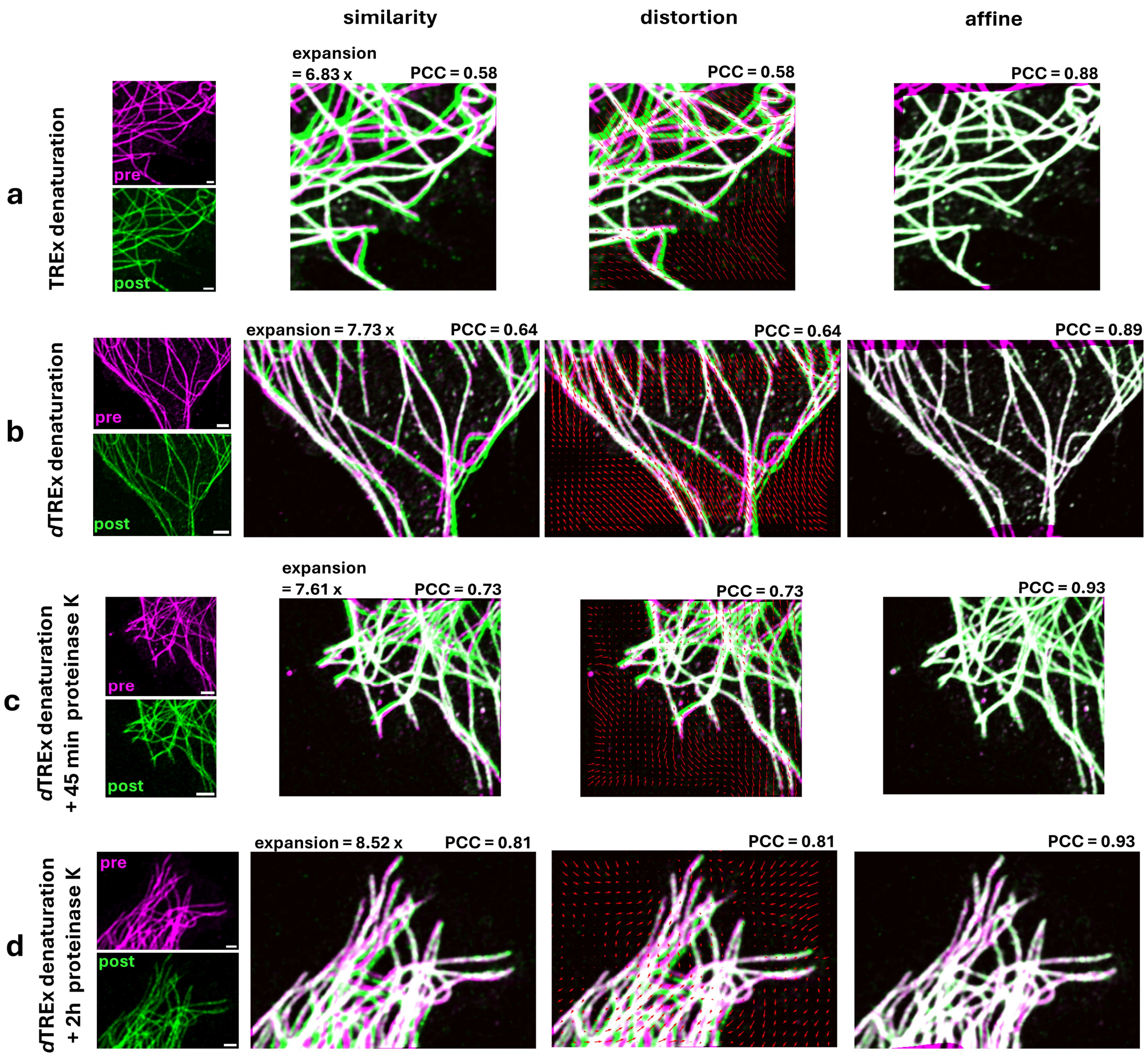


**Supplementary Fig. S2. Expansion factors of different protocols in COS-7 cells stained for alpha-tubulin.** Cells were fixed and anchored with GA. Denaturation was done with SDS and DTT at 98°C. In (c) and (d) proteinase K was applied at 37°C. Airyscan fluorescence images of the same area imaged pre (magenta) and post (green) expansion. Similarity transformation aligns pre- and post-expansion images by rotation, scaling and translation in x and y direction, yielding an expansion factor and PCC value. The distortion vector map was generated from the differences between similarity and non-rigid affine transformation, which usually yields a higher PCC. **(a)** TREx with denaturation. Scale bars pre: 1 µm, post: 10 µm. **(b)** *d*TREx with denaturation. Scale bars pre: 2 µm, post: 20 µm. **(c)** *d*TREx with denaturation and 45 min proteinase K. Scale bars pre: 2 µm, post: 20 µm. **(d)** *d*TREx with denaturation and 2 h proteinase K. Scale bars, pre: 1 µm, post: 10 µm. Scale bars show expanded dimensions.

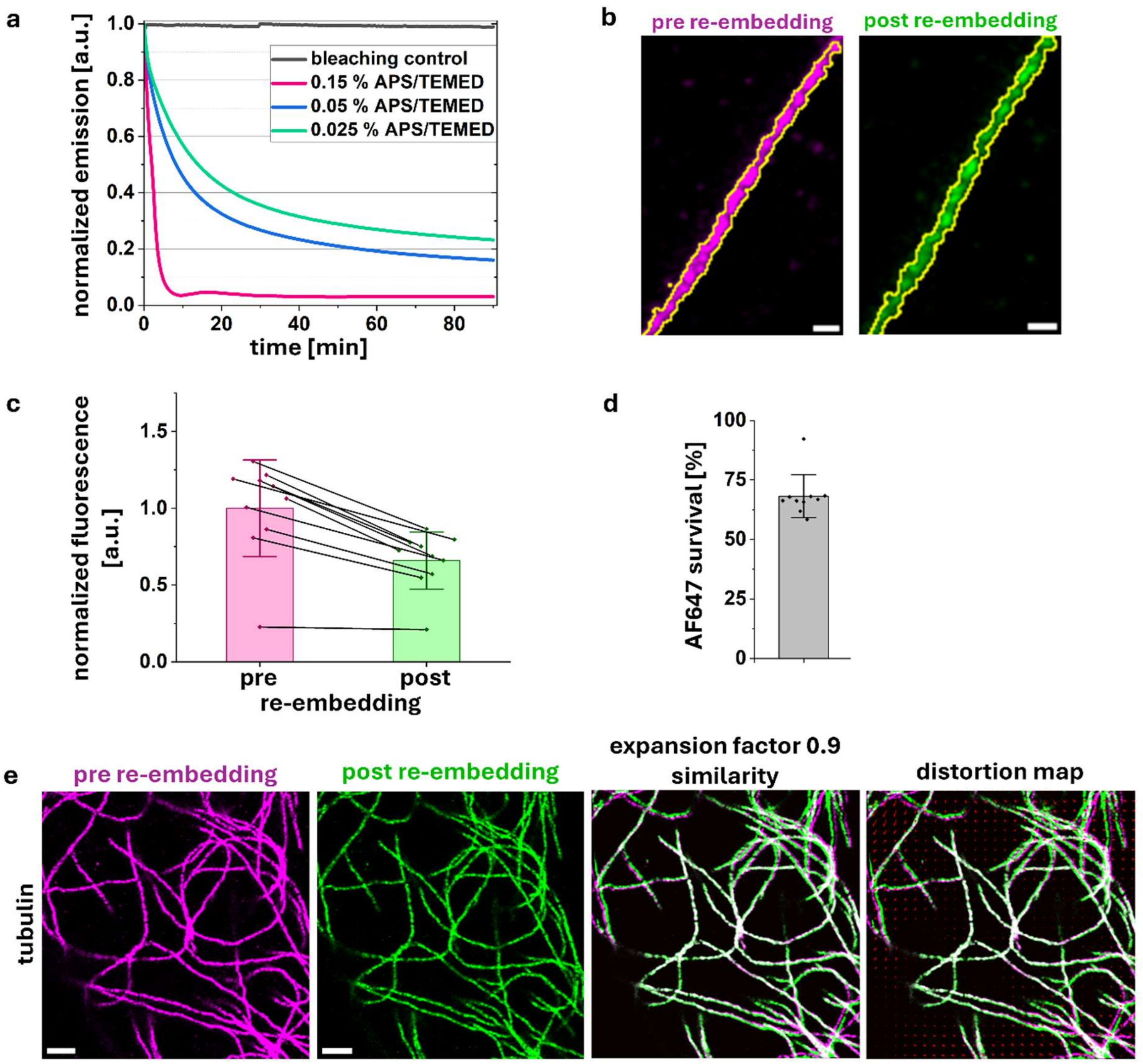


**Supplementary Fig. S3. AF647 survival and gel shrinkage during re-embedding in a neutral gel. (a)** AF647 ensemble emission intensity upon addition of APS and TEMED to an AF647-labeled antibody solution. After 90 min the normalized emission decreased to ~3 %, ~16 % and ~23 % using concentrations of 0.15 %, 0.05 % and 0.025 % APS/TEMED respectively. **(b)** Representative Airyscan images of an identical microtubule filament stained with AF647 coupled secondary antibody and imaged before (magenta) and after re-embedding (green). Yellow lines indicate ROI for fluorescence intensity measurement. **(c)** AF647 fluorescence intensity of tubulin ROIs (n = 10) normalized to the mean pre-re-embedding intensity. Lines indicate the fluorescence intensity loss for single data points. Data obtained from repeated measurements of one sample. **(d)** Mean AF647 fluorescence survival (68 ± 9 %). **(e)** Gel shrinkage during re-embedding. Identical areas of alpha-tubulin in COS-7 cells imaged pre (magenta) and post re-embedding (green). Images registered via similarity transformation yield an expansion factor of 0.9. A distortion vector map shows the difference to an affine transformation. Bar graphs show mean value ± s.d. Scale bars, 1 µm (B), 5 µm (E). Scale bars show expanded dimensions.

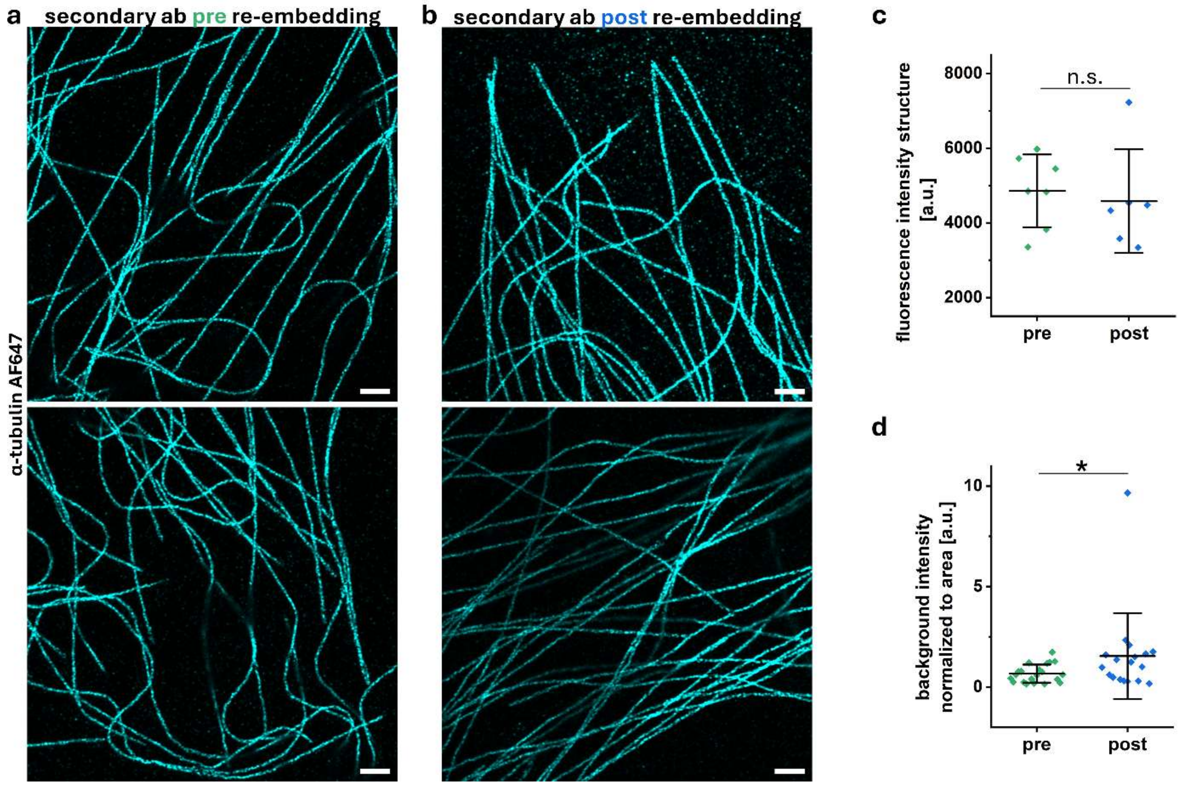


**Supplementary Fig. S4. Comparison of pre- and post-re-embedding labeling with secondary antibody. (a,b)** Representative Airyscan images of *d*TREx gels immunostained for α-tubulin and labeled with AF647-conjugated secondary antibodies either before (a) or after (b) re-embedding. **(c)** Images of microtubules labeled either before or after re-embedding with the secondary antibody show similar fluorescence intensities ($p = 0.68$ determined by two-sample t-test). **(d)** Background signal intensity of microtubule images labeled after re-embedding with the secondary antibody is slightly higher ($p = 0.047$ determined by Mann-Whitney-U test). Scale bars, 5 µm. Scale bars show 8.0x expanded dimensions after re-embedding in the neutral hydrogel.

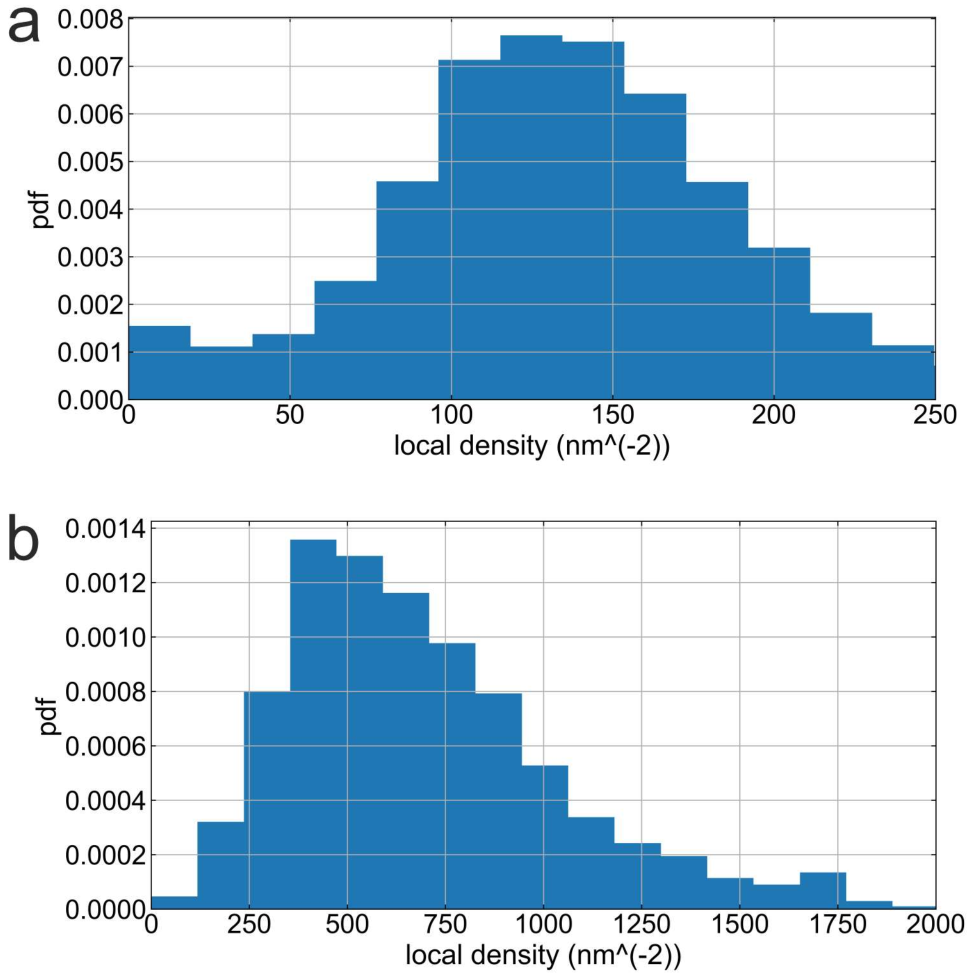


**Supplementary Fig. S5. Local densities of *d*STORM localizations for a-tubulin.** *d*STORM localizations in successive frames were linked when closer than 100 nm and the local density was computed for each linked localization with the given radius. **(a)** Histogram of localization densities (probability density function, pdf) detected for unexpanded microtubules using a 50 nm radius (n=23 ROIs). **(b)** Histogram of localization densities detected for ~8-fold expanded microtubules immunolabeled according to the *d*TREx protocol using a corresponding 400 nm radius (n=65 ROIs).

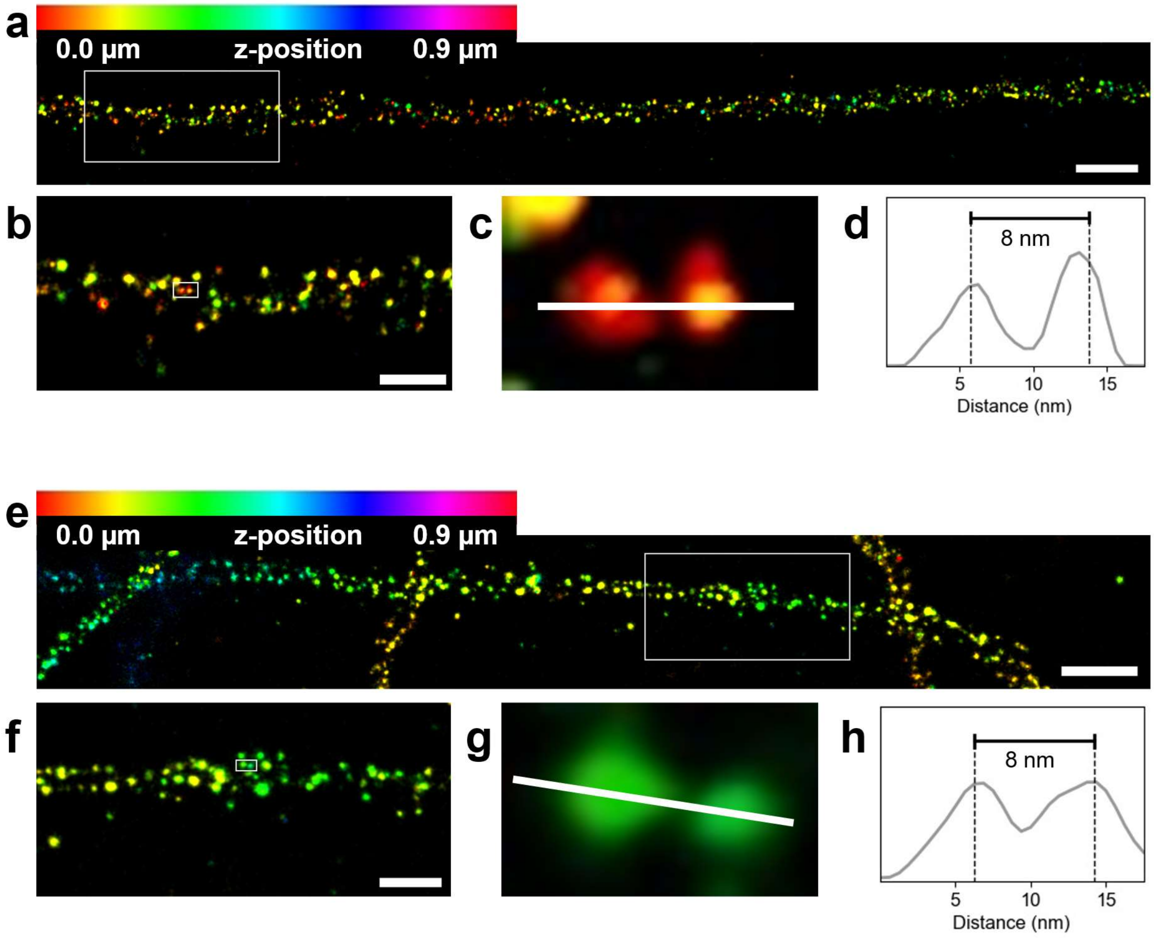


**Supplementary Fig. S6. Examples for 8 nm distances between tubulin dimers in cells resolved by Ex-*d*STORM. (a,e)** Microtubule segments in COS-7 cells imaged using *d*TREx post-immunolabeled for α-tubulin (AF647). **(b,f)** Zoomed-in views of regions indicated by the white box in (a) and (e). **(c,g)** Zoomed-in view of region indicated by the white box in (b) and (f). **(d,h)** Intensity profiles along the dotted lines in (c) and (g), with distances corrected for the final expansion factor of 8.0 after *d*TREx and re-embedding. Scale bars (expanded), a,e, 1 µm; b,f, 300 nm. Pixel size, 5 nm.

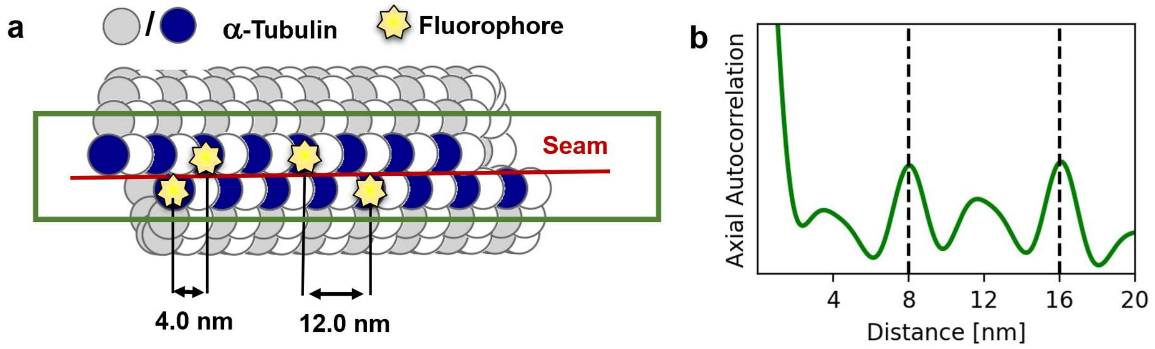


**Supplementary Fig. S7. Microtubule lattice seam in 13-1 configuration can lead to 4 nm periodicity in the axial autocorrelation. (a)** Schematic of simulated microtubule with α-tubulin indicated in blue/gray, with A-lattice contacts along the seam and measured distances between fluorophores that would result when analyzing the region indicated by the green box. (**b)** Axial autocorrelation along the seam of a simulated microtubule, analyzed with the same parameters as the experimental data.

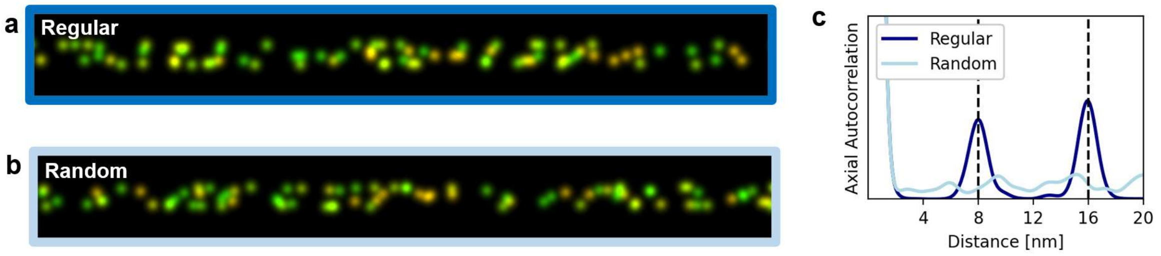


**Supplementary Fig. S8. Axial autocorrelation reveals the periodicity of the tubulin lattice.** **(a)** Simulation of an 8x expanded microtubule segment shown as color-coded z projection, with 5% of α-tubulin labelled, using the parameters extracted from experimental data. (**b)** Same data as in (a) with same number of fluorophores, but randomized positions and no underlying lattice periodicity. (**c**) Axial autocorrelation of regular (dark blue) and randomized (light blue) tubulin lattice in (a) and (b) reveals the underlying periodicity.

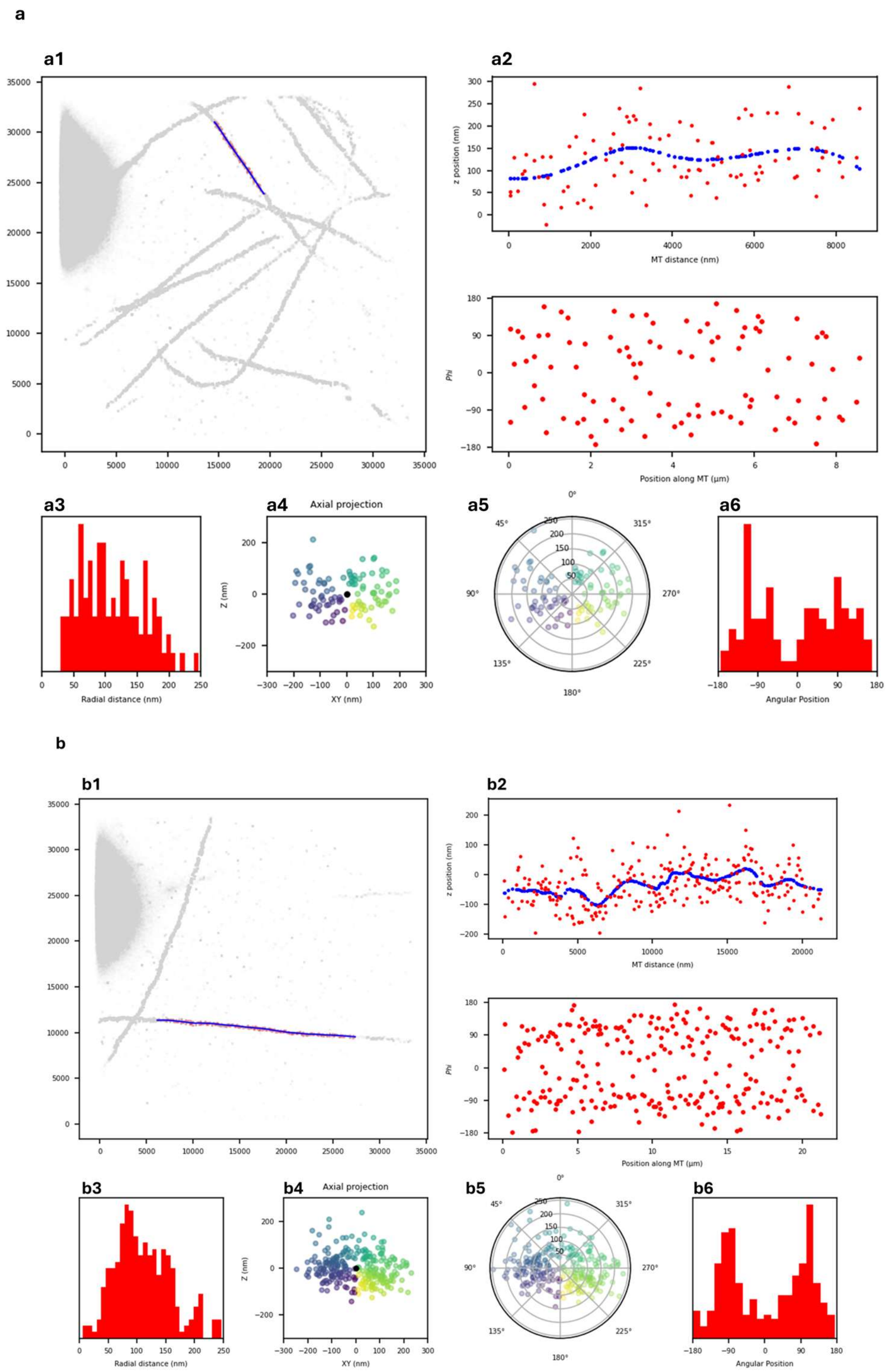


**Supplementary Fig. S9. Examples of analyzed regions of interest from Ex-*d*STORM images of microtubules. (a1,b1)** The blue line indicates the selected microtubule strand. **(a2,b2)** Z-position plotted over microtubule distance. The blue line shows the 3D centerline of cluster coordinates (red dots). **(a3,b3)** Angular positions of localization clusters (red dots) over microtubule distance. **(a3,b3)** Radial distance distribution of localization clusters. **(a4,b4)** Axial projection of localization clusters. **(a5,b5)** Angular positions of localization clusters as axial projection. **(a6,b6)** Distribution of angular positions.

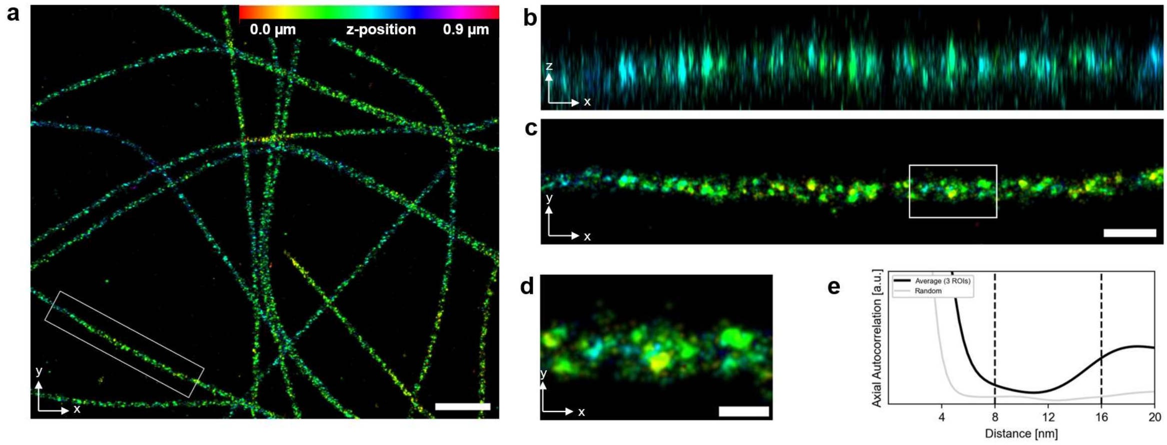


**Supplementary Fig. S10. Ex-*d*STORM of 3.2x expanded microtubules does not resolve the 8 nm tubulin lattice. (a)** Representative 3D Ex-*d*STORM image of re-embedded COS-7 cells expanded ~3.2-fold from Zwettler *et al*. (*66*). **b-c,** Corresponding xz- and xy-views of the region marked in (a). **(d)** Zoomed-in view of the region marked in (c). **(e)** Autocorrelation function averaged over three microtubule segments (black) and for the same number of clusters randomly placed across the filament (grey). Distance (x-axis) corrected for the expansion factor of 3.2x. Scale bars (expanded), a, 2 µm; b,c, 500 nm; d, 300 nm. Pixel size, 5 nm.

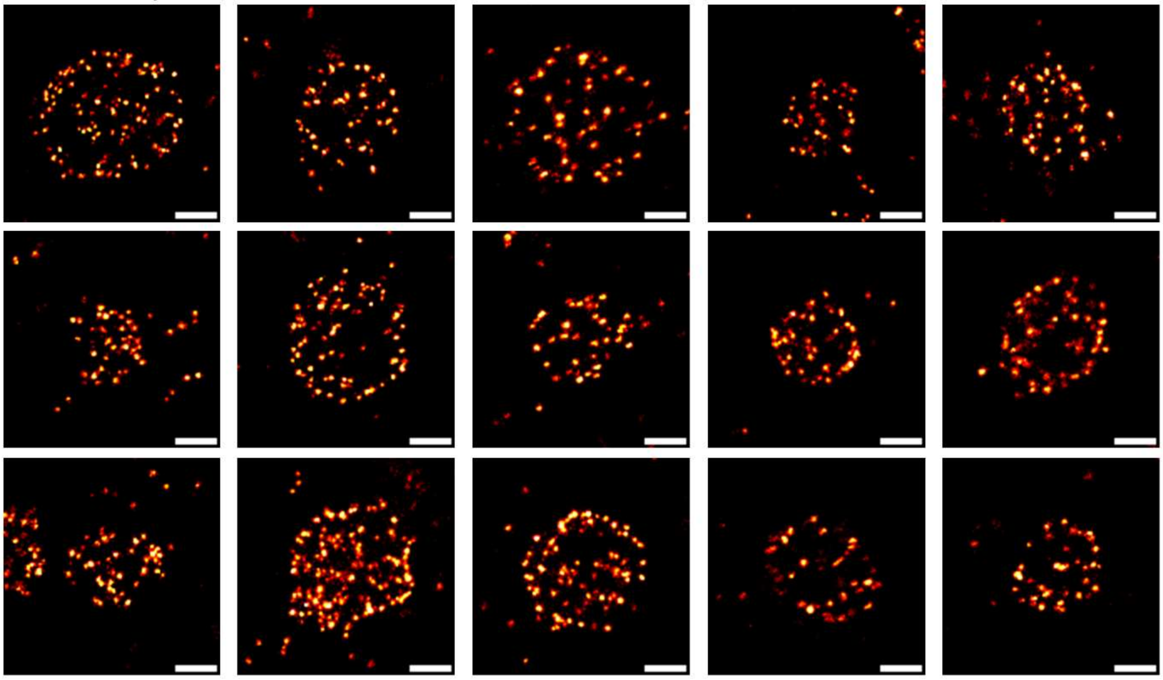


**Supplementary Fig. S11. Representative Ex-*d*STORM images of individual CCPs.** Gels of GA fixed and anchored COS-7 cells were immunostained for clathrin heavy chain after denaturation with 98°C. Samples were then processed according to *d*TREx including 45 min proteinase K digestion at 37°C. Results from three independent experiments. Scale bars 500 nm (7x expanded dimensions).

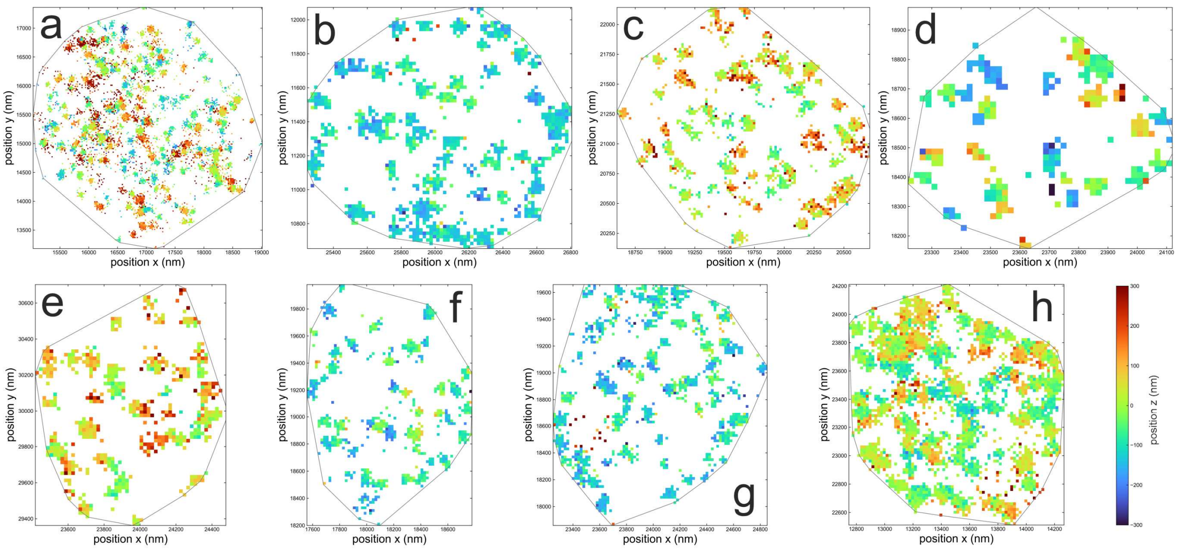


**Supplementary Fig. S12. CCPs in 3D-Ex- *d*STORM**. Data is shown for the regions of interest that were selected for radial distance distribution analysis in Supplementary Fig. 9. Individual localizations are binned in 20 nm pixels. The turbo color map represents the *z*-coordinate (average of all localizations per pixel). The gray region represents the 2D convex hull of all localizations projected in the xy-plane.

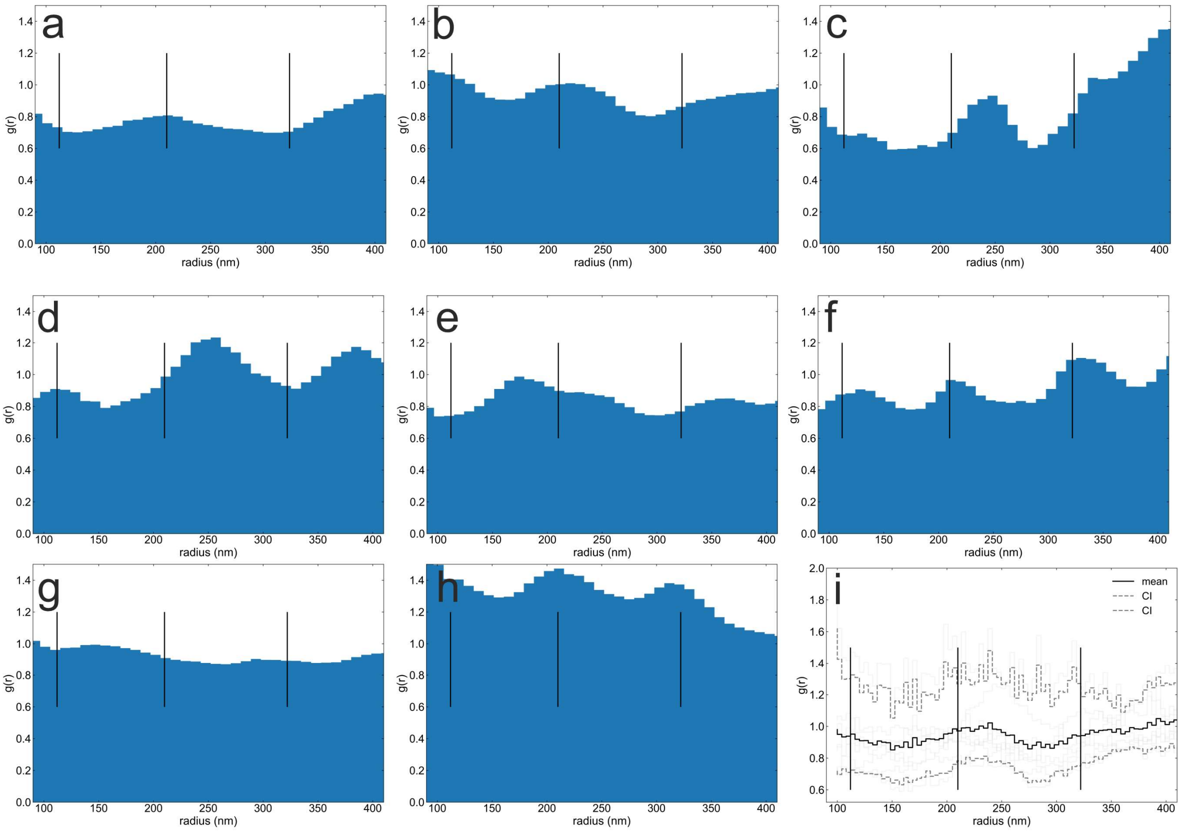


**Supplementary Fig. S13. Radial distribution function for individual CCP ROIs. (a-h)** The radial distribution function is shown for all pairwise localization distances from the recorded *d*STORM localization data. **(i)** The average radial distribution function with 5/95% confidence intervals for the same rois (same as in Fig. 3g). The vertical lines show expected peak positions. Radial distribution functions are shown relative to those for localizations distributed under complete spatial randomness in identical regions.

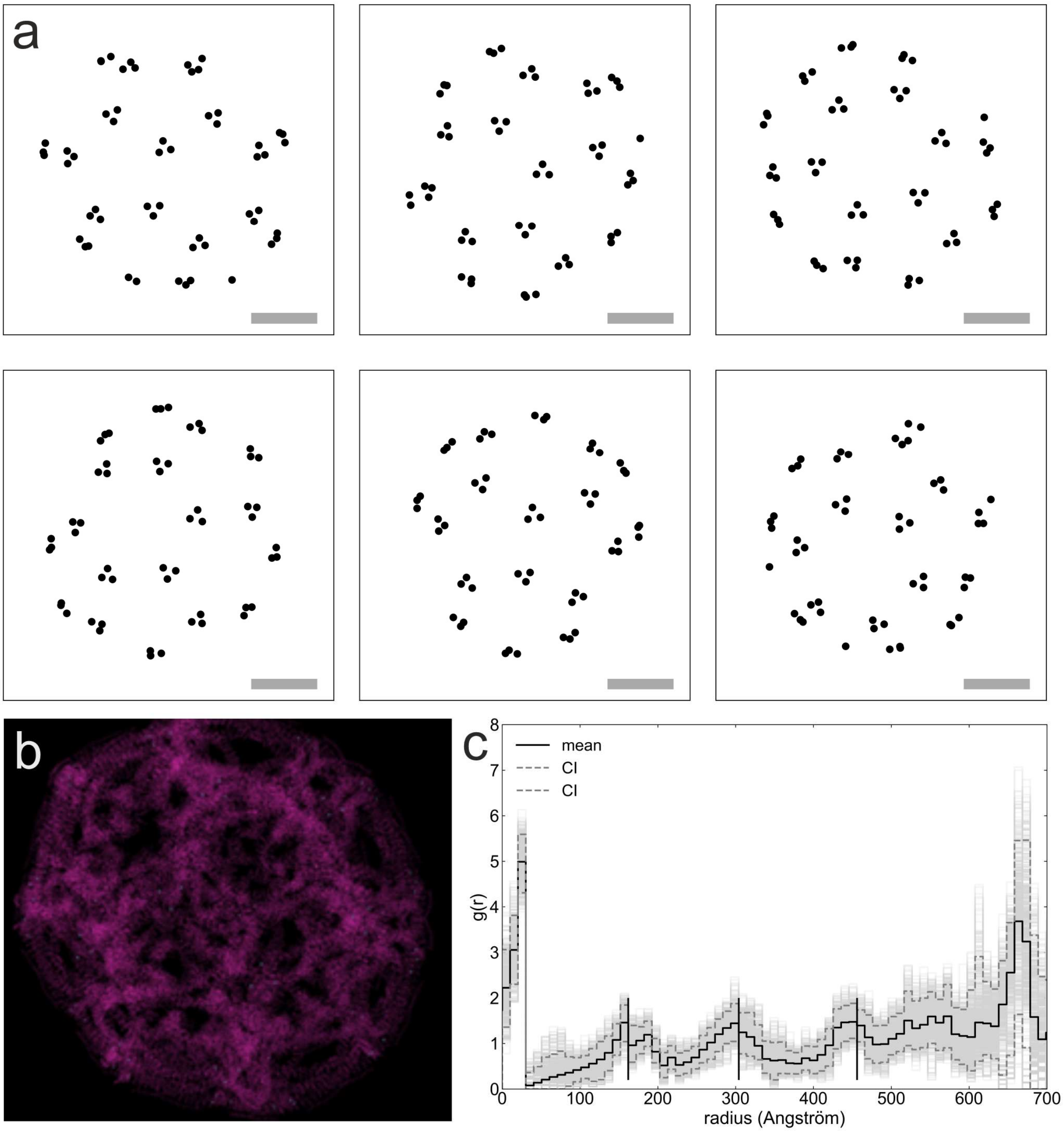


**Supplementary Fig. S14. Radial distribution functions for antibody epitope positions in CCP model structure. (a,b)** Vertices that represent the antibody epitopes on clathrin heavy chain (represented by amino acid 1551) are generated from pdb structure of clathrin D6 coat (pdb: 1XI4). For comparison with experimental data, the vertices on the upper half are selected and randomly rotated in 3D and then projected on the xy-plane. Scale bar: 20 nm. **(c)** Radial distribution functions for paired distances are generated and plotted (light gray lines) together with mean (black line) and 5/95 % confidence intervals (dashed gray lines) (n=100). Radial distribution functions are shown relative to a 2D spatial distribution under complete spatial randomness within the convex hull of all points combined.

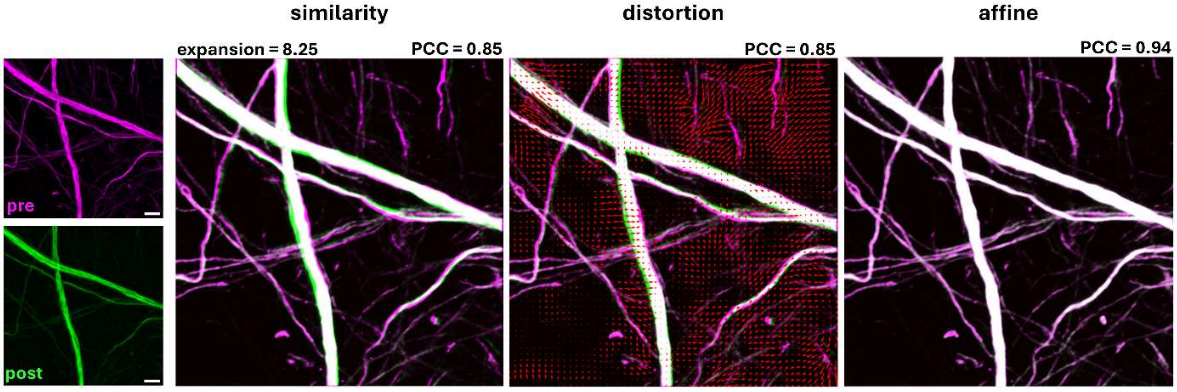


**Supplementary Fig. S15. Expansion factor of *d*TREx in neurons.** Primary hippocampal mouse neurons were fixed with FA and anchored with FA+AA. Neurofilament-H immunostaining was used to determine the expansion factor of *d*TREx using denaturation at 98°C and 45 min proteinase K digestion at 37°C. Airyscan images of the same area pre-expansion (magenta) and post-expansion (green) were registered by similarity transformation yielding an expansion factor of 8.25x, a PCC value and an affine transformation. The distortion vector map illustrates the differences between similarity and affine transformation. Considering ~10 % shrinking during re-embedding into the neutral gel, the expansion factor was estimated to ~7.5x in Ex-*d*STORM images. Scale bars, 3 µm (pre-expansion), 25 µm (post-expansion).

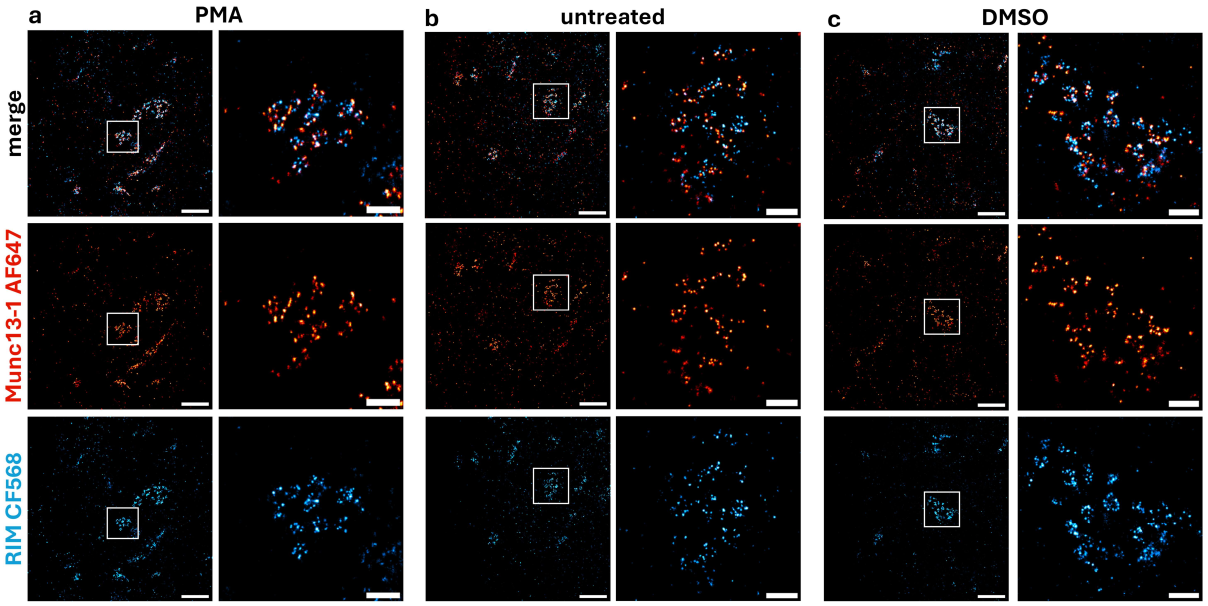


**Supplementary Fig. S16. Two-color Ex-*d*STORM images of Munc13-1 and RIM show frontal views of active zones of presynapses.** Hippocampal mouse neurons with different treatments before fixation with FA and anchoring with FA/AA, processed by *d*TREx using denaturation with SDS and DTT at 98°C and 45 min proteinase K at 37°C. From overview images a frontal view synapse was selected (white square) and magnified. **(a)** Neurons treated with PMA. **(b)** Untreated neurons. **(c)** Control with DMSO (solvent used for PMA treatment). Overview images scale bars, 5 µm, pixel size, 60 nm; magnified regions scale bars, 1 µm, pixel size, 20 nm. Scale bars show 7.5x expanded dimensions after re-embedding in the neutral hydrogel.

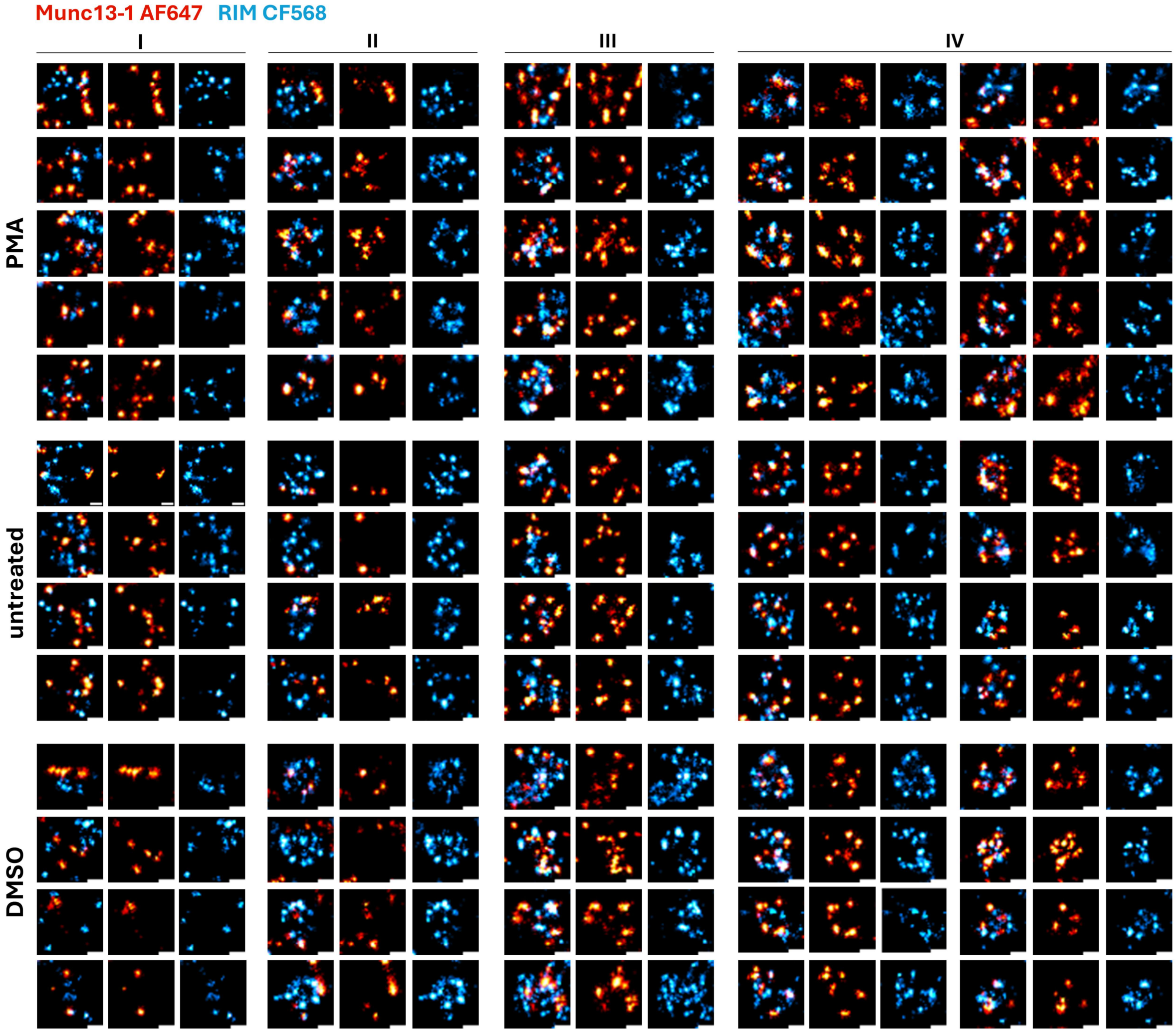


**Supplementary Fig. S17.** Hand-picked classification of ring-like structures of Munc13-1 and RIM into four classes under different experimental conditions. Magnified Ex-*d*STORM images of individual docking sites show regions of varying sub-structures from different synapses categorized in four different states. I: Munc13-1 and RIM unorganized. II: Only RIM shows ring-like arrangements. III: Munc13-1 and RIM are organized in substructures with a diameter > 500 nm. IV: Munc13-1 and RIM are both organized in ring-like structures with varying diameters. Scale bars, 200nm. Scale bars show 7.5x expanded dimensions after re-embedding in the neutral hydrogel.

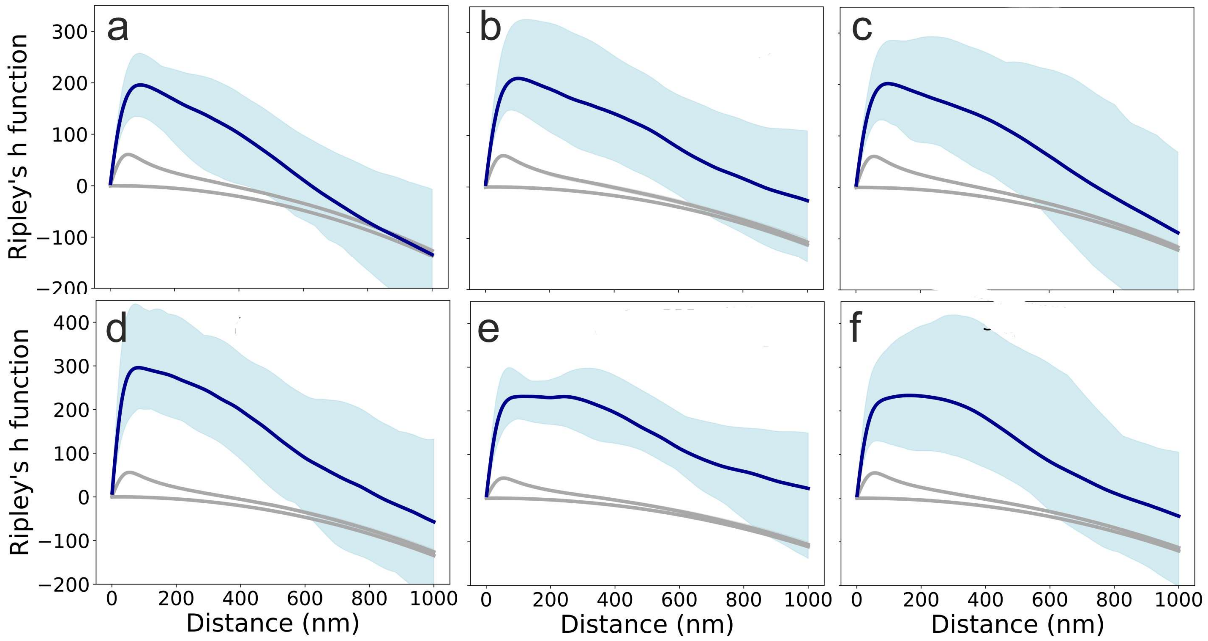


**Supplementary Fig. S18. Ripley h-function of *d*STORM localizations recorded for Munc13-1 and RIM immunolabeling.** The h-functions for experimental data are shown with a 5-95% confidence interval from multiple synapses for Munc13-1 (a-c) and RIM (d-e). The distance refers to expanded samples. Neurons were treated according to the following groups: untreated (a, d), DMSO-control (b,e), stimulated (c,f).The h-function from simulated data spatially distributed according to complete spatial randomness or according to a clustering process that resembles *d*STORM with homogeneously distributed emitters, is outside the experimental confidence interval for length scales up to ~100 nm (~750 nm expanded). This indicates clustering processes distinct from repetitive *d*STORM blinking on different clustering length scales. Scale bars represent ~7.5x expanded dimensions.

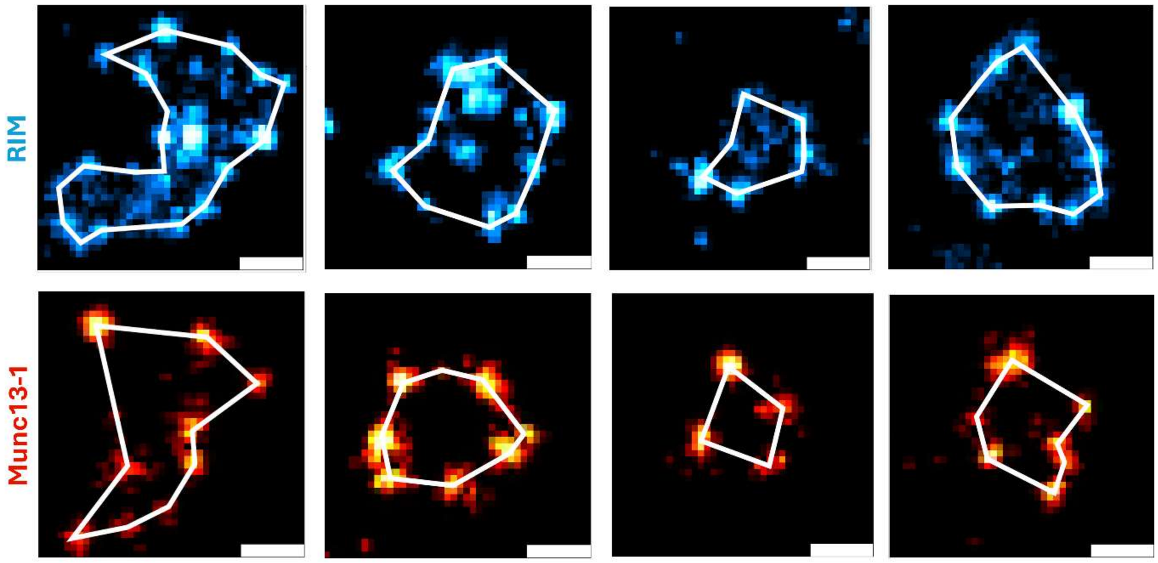


**Supplementary Fig. S19**. **Size analysis of Munc13-1 and RIM structures in individual synapses.** Synapses and substructures were identified as regions of interest by user selection. Using a polygon tool the outer signals of the respective structure were connected and the Feret's diameter was measured to determine the maximum diameter of the structure. This was done for selected structures of state II (only RIM), state III and state IV shown in Fig. 5b, fig. S17, and additional similar structures. Scale bars, 200 nm (7.5x expanded dimensions).

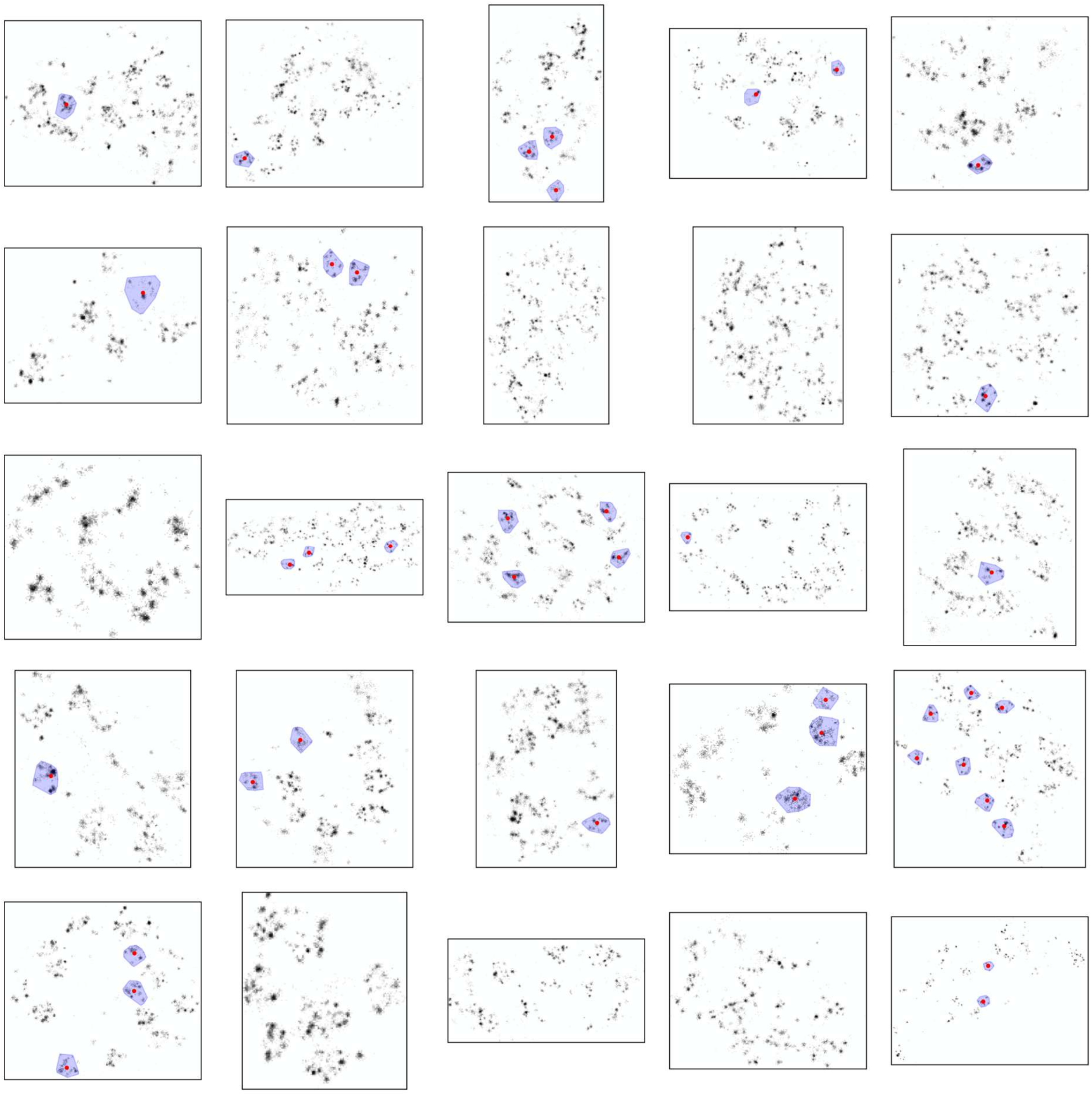

**Supplementary Fig. S20**. **Cluster identification of Munc13-1 and RIM signals in individual synapses.** Synapses were identified as regions of interest by user selection. DBSCAN was used to identify clusters of the combined set of Munc13-1 and RIM localizations within each synapse. A reproducible set of clusters was selected based on convex hull areas, circularity as represented by the isoperimetric ratio, and the radial distance for each cluster. Localization density is shown in gray. For all selected clusters, the convex hull region is shown in light blue and the centroid in red. It must be noted that the cluster selection does not represent a specific kind of cluster but only serves as objective identification procedure for a heterogeneous cluster set that contains larger ring-like structures. Scaling various throughout the panels; bin size = 20 nm.

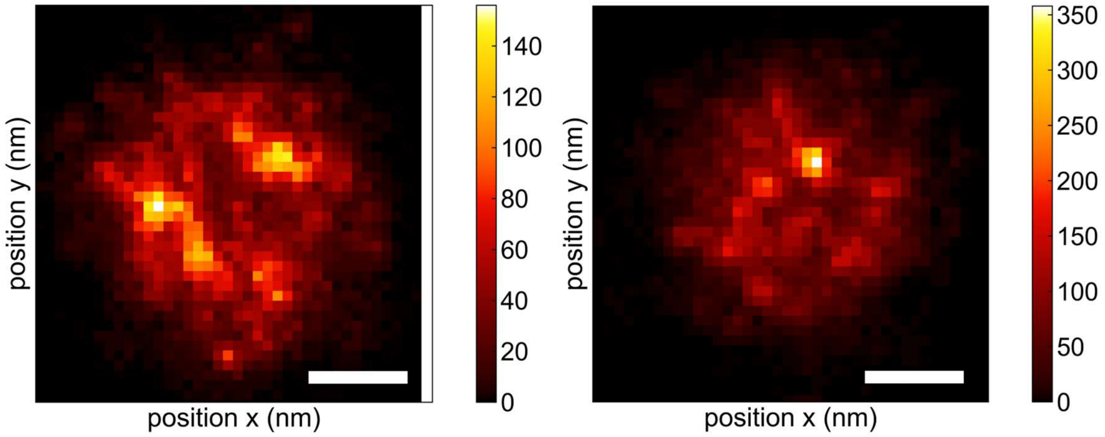


**Supplementary Fig. S21. Overlay images of localization clusters for Munc13-1 and RIM signals.** Clusters were determined on the combined Munc13-1 and RIM signals by DBSCAN and selected based on convex hull area and circularity (as described and shown in Fig. S6), shifted by their centroid position and rebinned (bin size=20 nm) as overlay figure. The spatial distribution indicates the variety of substructures hiding the center hole that clearly appears in selected clusters. Color scale represents localizations per pixel. Scale bar, 200 nm (7.5x expanded dimensions).

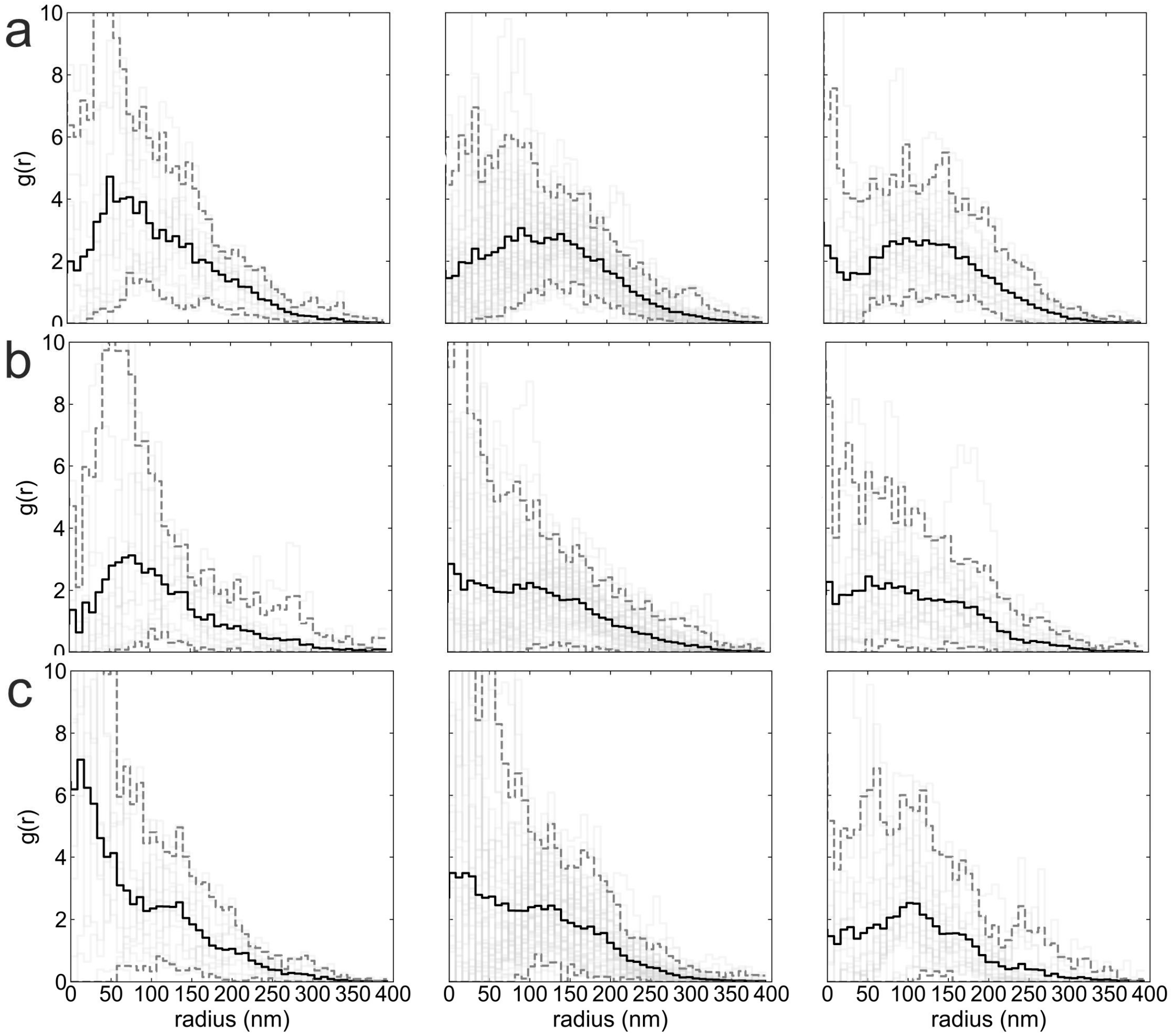


**Supplementary Fig. S22. Radial distribution function for clusters of Munc13-1 and RIM signals.** The radial distribution function is shown for all distances between localizations in Munc13-1 and RIM clusters and the cluster centroid in expanded samples: **(a)** The radial distribution function for the combined Munc13-1 and RIM signals. **(b)** The radial distribution function for Munc13-1 signals. **(c)** The radial distribution function for RIM signals. In all panels data is shown for treatments DMSO-control (left), stimulated (center) and untreated (right). Radial distribution functions are plotted (light gray lines) together with mean (black line) and 5/95 % confidence intervals (dashed gray lines). Scales represent 7.5x expanded dimensions.

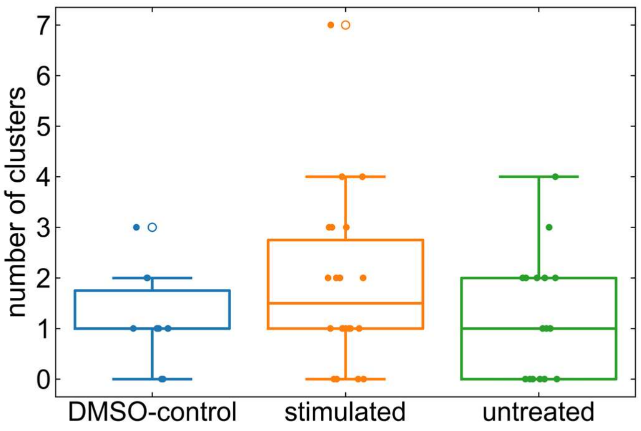


**Supplementary Fig. S23. Number of clusters of Munc13-1 and RIM signals per region of interest.** Clusters were determined as described and shown in Supplementary Fig. 13. The mean number of clusters were not significantly different between the various treatments.

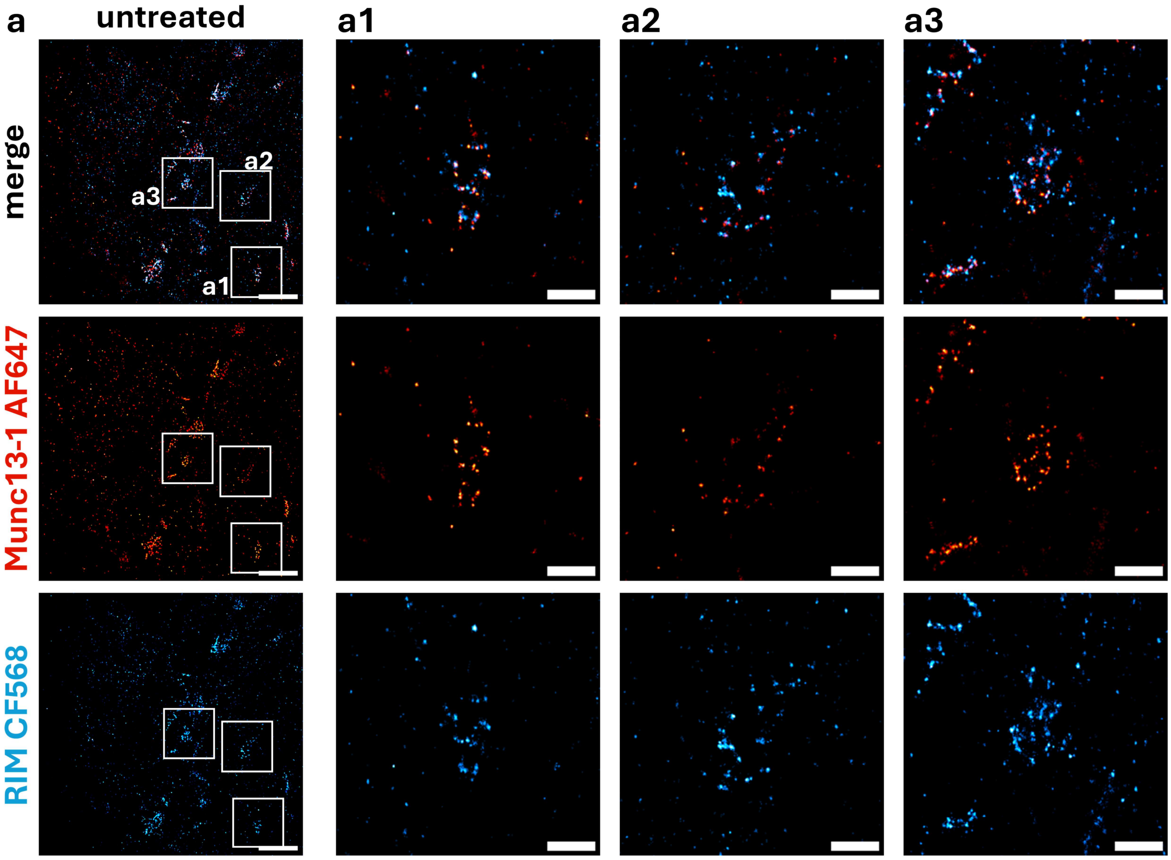


**Supplementary Fig. S24. Two-color Ex-*d*STORM images of Munc13-1 and RIM1/2 using only denaturation during expansion.** Untreated hippocampal mouse neurons were processed with a single TREx hydrogel using FA fixation, FA/AA anchoring and denaturation with SDS and DTT at 98°C and no proteinase K digestion. Substructures lack details and seem not properly expanded. **(a)** Representative overview image. Selected frontal views of synapses are marked by a white square and magnified in a1, a2 and a3. Scale bars, a, 5 µm; a1-a3, 1 µm. Pixel size a, 60 nm; a1-a3, 20 nm. Scale bars show expanded dimensions with an estimated expansion factor of 5-6 (single TREx gel shrinks ~20 % during re-embedding in the neutral hydrogel).
